# Improved estimation of macroevolutionary rates from fossil data using a Bayesian framework

**DOI:** 10.1101/316992

**Authors:** Daniele Silvestro, Alexandre Antonelli, Nicolas Salamin, Xavier Meyer

**Affiliations:** Department of Biological and Environmental Sciences, University of Gothenburg, 413 19 Gothenburg, Sweden; Global Gothenburg Biodiversity Center, Gothenburg, Sweden; Department of Computational Biology, University of Lausanne, 1015 Lausanne, Switzerland; Swiss Institute of Bioinformatics, Quartier Sorge, 1015 Lausanne, Switzerland; Gothenburg Botanical Garden, SE-41319 Goteborg, Sweden; Department of Organismic and Evolutionary Biology, Harvard University, 26 Oxford St., Cambridge, MA 02138 USA; Department of Integrative Biology, University of California, Berkeley, CA 94720, USA

**Keywords:** PyRate, origination and extinction rates, Reversible Jump MCMC, birth-death models

## Abstract

The estimation of origination and extinction rates and their temporal variation is central to understanding diversity patterns and the evolutionary history of clades. The fossil record provides the most direct evidence of extinction and biodiversity changes through time and has long been used to infer the dynamics of diversity changes in deep time. The software PyRate implements a Bayesian framework to analyze fossil occurrence data to estimate the rates of preservation, origination and extinction while incorporating several sources of uncertainty. This fully probabilistic approach allows us to explicitly assess the statistical support of alternative macroevolutionary hypotheses and to infer credible intervals around parameter estimates. Here, we present a major update of the software, which implements substantial methodological advancements, including more complex and realistic models of preservation, a reversible jump Markov chain Monte Carlo algorithm to estimate origination and extinction rates and their temporal variation, and a substantial boost in performance. We demonstrate the new functionalities through extensive simulations and with the analysis of a large dataset of Cenozoic marine mammals. We identify several significant shifts in origination and extinction rates of marine mammals, underlying a late Miocene diversity peak and a subsequent 50% diversity decline towards the present. Our analyses indicate that explicit statistical model testing, which is often neglected in fossil-based macroevolutionary analyses, is crucial to obtain accurate and robust results. PyRate provides a flexible, statistically sound analytical framework, which we think can serve as a useful toolkit for many future studies in paleobiology.

## Introduction

The evolution of biological diversity is determined by the interplay between origination and extinction processes. Estimating the pace at which lineages appear and disappear is therefore a central question in macroevolution and paleobiology research. Inferring the processes underlying biodiversity patterns helps us understanding what drives the wax and wane of taxa (Ezard et al., 2011; Quental and Marshall, 2013), the effects of competition and other biotic interactions on diversity changes (Liow et al., 2015; Pires et al., 2017), the dynamics and selectivity of mass extinctions (Peters, 2008). The process of taxonomic diversification is often modeled using birth-death stochastic models, where the appearance of new lineages (e.g. species or genera) and their demise are characterized by origination and extinction rates (Kendall, 1948; Keiding, 1975; Nee, 2006). These parameters quantify the expected number of origination or extinction events per lineage per time unit (typically 1 million years) (Foote, 2000; Marshall, 2017).

In recent years, there have been considerable methodological developments in the estimation of diversification dynamics from phylogenies of extant taxa, in which the distribution of branching times calibrated to absolute ages are used to infer the parameters of a “reconstructed birth-death process” (e.g. Nee et al., 1994; Gernhard, 2008; Stadler, 2009, 2013; Heath et al., 2014). These methods are appealing because large phylogenies of extant taxa are becoming increasingly available (e.g. Jetz et al., 2012; Pyron et al., 2013; Zanne et al., 2014; Rolland et al., 2018) and extend to taxa with limited fossil record, including hyper-diverse clades such as orchids (Perez-Escobar et al., 2017). Despite this methodological progress, there are limitations to estimating diversification dynamics from extant data, particularly in terms of estimating realistic extinction rates (Rabosky, 2010; Quental and Marshall, 2010; Liow et al., 2010a; Marshall, 2017). A major limiting factor of phylogenetic approaches to infer origination and extinction rates is that extant species represent, for most clades, a small fraction of a the total diversity that has existed since their origination (Raup and Sepkoski, 1984; Raup, 1986).

The fossil record provides the most direct evidence of past biodiversity and extinction and has therefore long been used to investigate diversification processes (Kurtén, 1954; Van Valen and E, 1966; Alroy, 1996; Sepkoski, 1998; Alroy, 2008; Foote, 2001; Liow and Nichols, 2010; Ezard et al., 2011). However, since the paleontological record is virtually always incomplete, fossil occurrences represent a biased representation of the past diversity, where the sampled longevities of taxa are likely to underestimate their true lifespan, and entire lineages (especially those with low preservation potential or short lifespan) may leave no trace of their existence (Foote, 2000; Foote and Raup, 1996; Hagen et al., 2017). Thus, the estimation of diversification processes from fossil data typically involves inferring preservation, origination, and extinction rates. Most available methods estimate temporal rate variation using the presence or absence of lineages within predefined time bins and treating the origination and extinction rates in each bin as independent parameters (Foote, 2001, 2003; Liow et al., 2008; Liow and Nichols, 2010; Alroy, 2014). The resulting patterns usually depict rate fluctuations through time, which may however capture stochastic variations from a time-homogeneous birth-death process and potentially reflect the problems of overparameterization, i.e. overfitting associated with the use of a higher number of parameters than supported by the data (Burnham and Anderson, 2002).

A few years ago we presented a Bayesian probabilistic framework to estimate preservation, origination and extinction rates from fossil occurrence data implemented in the open-source program PyRate (Silvestro et al., 2014b, a). Unlike most other methods, PyRate does not by default estimate origination and extinction rates within fixed time bins (although it is able to do it, as shown in Silvestro et al., 2015b). Instead, its core functions are designed to explicitly compare models with different amounts of rate heterogeneity, with the rationale that rate shifts are only detected when statistically significant. This procedure is important to avoid overparameterization, which in turn can lead to inconsistent results and false positives. This is especially true when the amount of data is small compared to the number of parameters (Burnham and Anderson, 2002), which is often the case for empirical fossil datasets.

Since its original implementation, PyRate uses a hierarchical Bayesian model to jointly estimate: 1) the times of origination and extinction for each sampled lineage (Fig. 1A), 2) the parameters of a Poisson process modeling fossilization and sampling (Fig. 1B), 3) the rates of origination and extinction and their temporal heterogeneity (Fig. 1C) (Silvestro et al., 2014a). This hierarchical structure allows us to analyze the entire available fossil record including all known occurrences of a lineage (i.e. not limited to first and last appearances), singletons (lineages sampled in a single occurrence), and extant taxa provided that they have at least one fossil occurrence (Fig. 1A). The analysis is conducted using Metropolis Hastings Markov chain Monte Carlo (MCMC), to obtain posterior estimates of all model parameters along with the respective 95% credible intervals (95% CI), providing important information about the level of uncertainty surrounding the estimates. One of the main and most challenging aims of the PyRate method is the estimation of how origination and rates vary through time. In its initial implementation, PyRate included a birth-death MCMC (BDMCMC) algorithm (Stephens, 2000) to sample the number and temporal placement of rate shifts in a single analysis. The power of this algorithm, however, appears to become limited with increasing levels of rate heterogeneity through time and with large datasets (Silvestro et al., 2014b).

**Figure 1:**
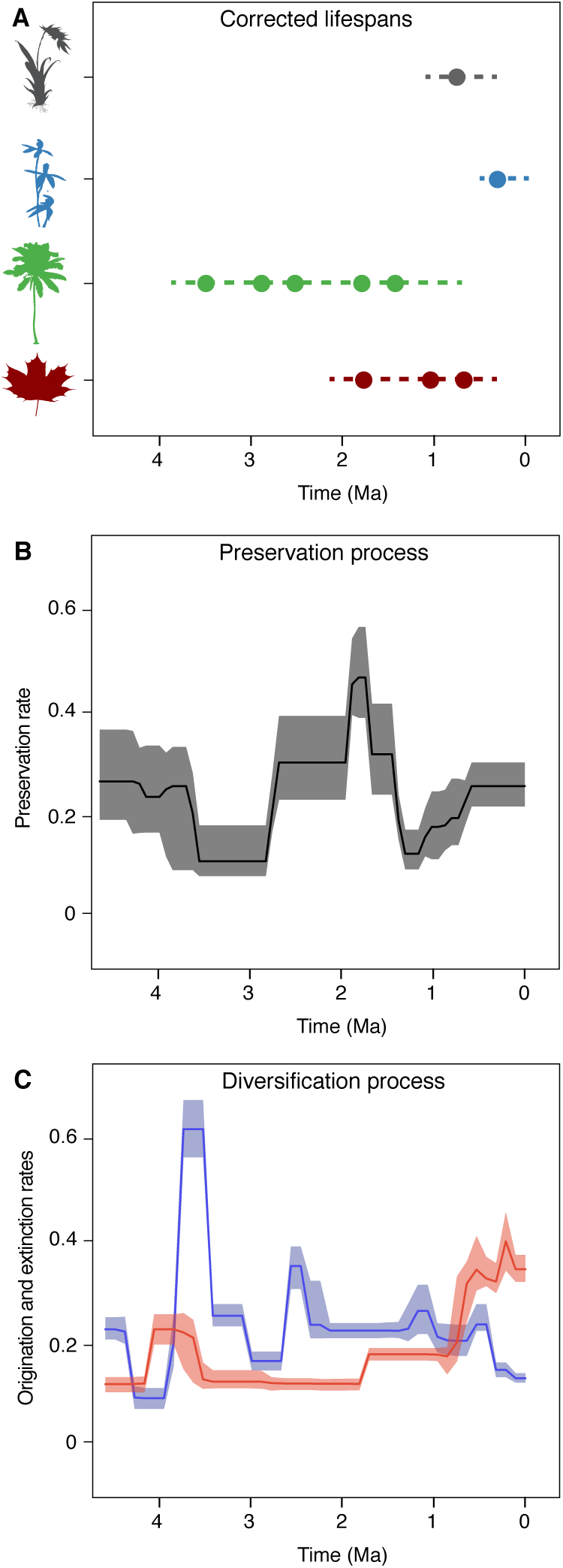
PyRate’s main analytical structure. The input data consist of dated fossil occurrences assigned to lineages, e.g. species or genera (represented by circles in A), including singletons and extant taxa. The Bayesian framework jointly estimates the lifespans of all lineages (dashed lines), preservation rates (B) and origination and extinction rates (C). All parameter estimates are inferred as posterior mean values (solid lines in B and C) and 95% credible intervals (shaded areas in B and C).

Here we develop extensive improvements of the PyRate method and present a substantially upgraded version of software introducing several novel features, which expand the scope and applicability of the program for the paleobiological community and improve user experience. Specifically we 1) introduce more realistic preservation models simultaneously allowing rate heterogeneity across lineages and through time. 2) We develop a new model testing framework using maximum likelihood to choose among alternative preservation models. 3) We present a more powerful algorithm to infer temporal variation in origination and extinction rates using reversible jump MCMC (RJMCMC) and compare its performance with the alternative BDMCMC algorithm, demonstrating improved results on simulated data. 4) We develop FastPyRateC, a C++ library which is seamlessly imported by the main PyRate program and yields a dramatic boost in performance, by optimizing the likelihood computations. FastPyRateC can speed up the analyses by orders of magnitude and the performance gain increases with the size of the dataset and the complexity of the model. 5) We provide a number of new functions to process output files and plot the results, calculate timing of significant rate shifts based on Bayes factors, and assess the presence of potential typos and misspellings in the taxa names in an input file. We demonstrate some of these features with a worked example by analyzing a recently published dataset of marine mammals (Pimiento et al., 2017) and provide extensive tutorials with detailed descriptions of analysis setup and output processing.

## Methods

PyRate implements a hierarchical Bayesian model that jointly samples the preservation rates (indicated by *q*), the times of origination and extinction for each sampled lineage (indicated by vectors **s**, **e**), and the origination and extinction rates (indicated by *λ* and *μ*). The input data are fossil occurrences characterized by their age and their assignment to a taxonomic unit (e.g. a genus or a species) and the origination and extinction rates scaled to the taxonomic unit utilized in the input data. The joint posterior distribution of all parameters is approximated by a Markov Chain Monte Carlo (MCMC) algorithm and can be written as

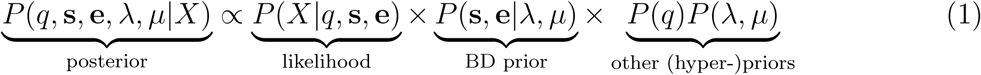

where *X* = {**x**_1_*,…***x**_*N*_ } is the list of vectors of fossil occurrences for each of *N* lineages, so that **x_i_** = {*x*_1_*,…, x_K_*} is a vector of all fossil occurrences sampled for taxon *i*. The likelihood component of the model allows us to estimate the preservation rates and the times of origin and extinction given the occurrence data, based on a stochastic model of fossilization and sampling (see below). The birth-death (BD) prior allows us to infer the underlying diversification process based on the (estimated) origination and extinction times. Additional priors on *q, λ, μ* enable the estimation of these parameters from the data. These priors are by default set to gamma distributions (thus allowing only positive values), unless otherwise specified.

### Preservation models

We model the process of fossil preservation and sampling using Poisson processes, where the estimated preservation rate(s) indicate the expected number of fossil occurrences per sampled lineage per time unit. Thus, fossil preservation is modeled as a time-continuous stochastic process capturing fossilization, sampling and identification, i.e. all the events occurring from the living organism to the digitized fossil occurrence. The likelihood of a lineage with fossil occurrences **x** = {*x*_1_*,…, x_K_*} given origination time *s*, extinction time *e*, and preservation rate *q* under a general Poisson model is

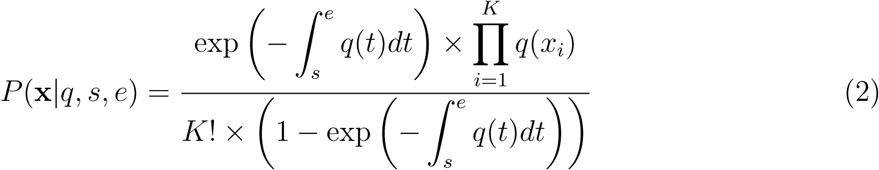

where *q*(*t*) is the preservation rate at time *t* (Silvestro et al., 2014b). The two terms of the numerator quantify the probability of the waiting times between fossil occurrences and the probability of each occurrence. The denominator includes the normalizing constant of the Poisson distribution and the condition on sampling at least one fossil occurrence, where exp(*⋅*) represents the probability of zero fossil occurrences between origination and extinction times (Silvestro et al., 2014b).

The original PyRate implementation included two models of preservation: the homogeneous Poisson process (HPP) and the non-homogeneous Poisson process (NHPP). The HPP model assumes that the preservation rate is constant throughout the lifespan of an organism and across time. The NHPP assumes that preservation rates change along the lifespan of a lineage according to a bell-shaped distribution, where the rates are lower at the two extremities (i.e., close to the times of origin and extinction of the lineage) and highest in the middle (Silvestro et al., 2014b). The shape of the distribution is fixed and the estimated preservation rate *q* represents the expected number of fossil occurrences per sampled lineage per Myr averaged across the lifespan of the lineage. This model is justified by the empirical observation that the number of occurrences per time unit for a given organisms tends to increase following its origination and to decrease prior to its extinction (Liow et al., 2010b). The pattern also reflects the idea that species originate from a small initial pool of individuals in a restricted geographic area (therefore with lower potential for preservation and sampling) and later expand, thus increasing the chances to leave fossil records. Similarly, under this model, species are expected to decline in abundance and geographic range prior to their extinction (Raia et al., 2016), resulting in decreased preservation rates.

Both HPP and NHPP models can be coupled with a Gamma model (i.e. HPP+G and NHPP+G), which allows us to incorporate rate heterogeneity across lineages. Under these models, preservation rates are defined so that their mean equals *q* and their heterogeneity is distributed according to a gamma distribution, with shape parameter *α*, discretized in a user-defined number of categories (Yang, 1994; Silvestro et al., 2014b). Both *q* and *α* are estimated as free parameters by the MCMC and small values of *α* indicate increased amount of heterogeneity. Gamma models do not assign individual preservation rates to each lineage in the dataset. Instead, the likelihood of each lineage is averaged across all rates, thus incorporating rate heterogeneity across lineages while adding a single additional parameter (*α*) to the model (Yang, 1994).

Here, we introduce a third preservation model, that implements a time-variable Poisson process (TPP). The TPP model is an extension of the HPP, in which the rate of preservation is constant within predefined time windows, but allowed to change between them. For instance, different preservation rates can be estimated within geological epochs (Foote, 2001; Liow and Nichols, 2010). The likelihood of this process is the product of piece-wise HPP likelihoods across multiple time frames, each with its specific preservation rate (**q** = {*q*_1_*,…, q_S_*}, where *S* is the number of time frames in the model). As for HPP and NHPP models, the TPP can be coupled with a Gamma model, therefore allowing for rate heterogeneity both through time and across lineages.

The default prior specified for *q* is a gamma distribution, chosen to reflect the fact that preservation rates must take positive values. Defining appropriate prior distributions is often a challenge in Bayesian analysis and prior choice can strongly affect the effective parameter space and the complexity of a model (Gelman et al., 2004). This may become even more problematic under the TPP model, where very strict priors could artificially reduce rate heterogeneity through time, whereas very vague priors could unnecessarily expand the amount of parameter space, increasing the risk of over-parameterization. To overcome this issue, we use a hyper-prior to estimate the prior on the preservation rates from the data, instead of setting the prior to a fixed distribution. We set a gamma prior on the vector **q** with fixed shape parameter (*α* = 1.5) and unknown rate parameter *β*. The rate parameter is assigned a vague gamma hyper-prior, *β ~* Γ(*a* = 1.01*, b* = 0.1), and is itself estimated from the data. Using the properties of the conjugate gamma prior, we sample the rate parameter *β* directly from its posterior distribution, given any vector of preservation rates **q**:

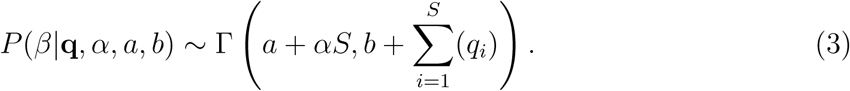

### A maximum likelihood test to compare preservation models

We developed a likelihood-based test to assess the statistical fit of alternative preservation processes. Although it is theoretically possible to infer the marginal likelihood of a preservation model in a Bayesian framework (for instance using the thermodynamic integration available in PyRate to test between alternative birth-death models (Lartillot and Philippe, 2006; Silvestro et al., 2014b)), the task would be computationally extremely demanding. Indeed, the number of parameters over which the likelihood needs to be marginalized can be very high, including the vectors of origination and extinction times, the preservation rates and potentially the parameters of the birth-death prior. Thus, we implemented a maximum likelihood test for preservation models, which substantially reduce computational burden.

Let *ŝ* and *ê* be the expected times of origination and extinction of a lineage with fossil occurrences **x** = {*x*_1_*,…, x_K_*} (sorted from oldest to most recent) for a given preservation rate *q*. In order to compare the fit of different models we maximize the likelihood *P* (**x***, ŝ, ê|q*), where *q* is treated as a free parameter and estimated in the optimization, while *ŝ* and *ê* are calculated based on the preservation rate and model. In the simplest case of an HPP of preservation the expected times of origination and extinction are determined by the expectation of an exponential distribution with rate equal *q*: **E**[*Exp*(*q*)] = 1*/q*. Thus, under HPP the expected times of origination and extinction are *ŝ* = *x*_1_ + 1*/q* and *ê* = *x_K_ −* 1*/q* (Fig. 2A). Note that the expected times of origination and extinction differ from their maximum likelihood estimates, which under HPP are *s_ML_* = *x*_1_ and *e_ML_* = *x_K_*.

**Figure 2:**
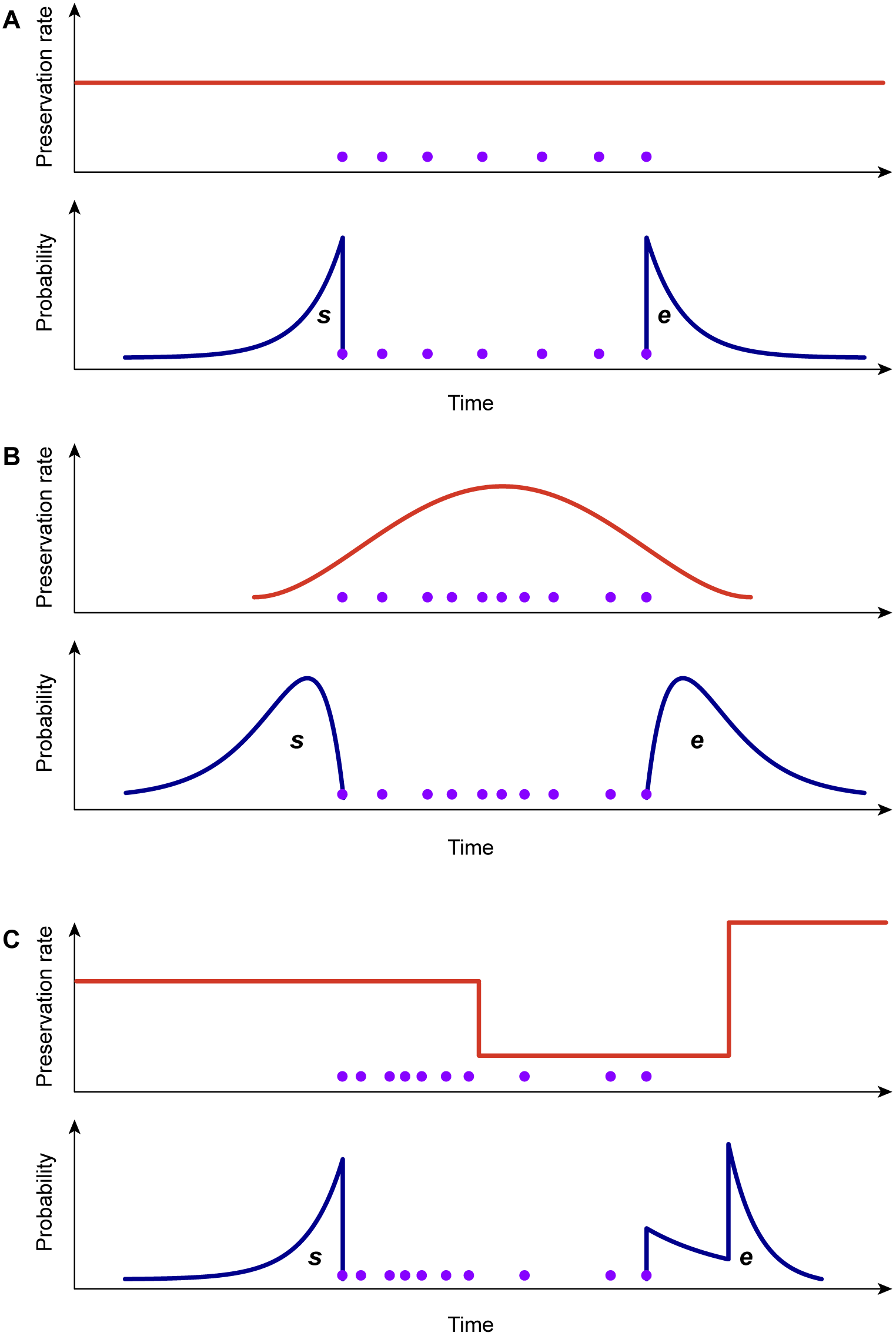
Graphical representation of the preservation rate models implemented in PyRate. In the HPP model (A) the preservation rate is constant through time and the expected times of origination and extinction (*s, e*, blue curves) are exponentially distributed. In the NHPP model (B), preservation rates vary throughout the lifespan of a species generating gamma-like expected *s, e*. The TPP model (C) assumes piece-wise constant preservation rates (e.g. different rates for each Epoch) and the resulting expected *s, e* combine multiple exponential distributions. All models can incorporate rate heterogeneity across-lineages (Gamma models).

In the case of the NHPP model, neither the expectation nor the maximum likelihood values of *s* and *e* are easily derived analytically. Instead, we use a two-step approach to obtain a maximum likelihood value that is comparable to that obtained under HPP. First, we optimize the rate *q* by maximizing the likelihood *P* (**x***|q, s, e*), where *q, s*, and *e* are treated as free parameters. This results in maximum likelihood estimates of the preservation rate *q_ML_* and origination and extinction times (*s_ML_* and *e_ML_*). Secondly, since the likelihoods of different preservation models are compared based on the expected origination and extinction times (i.e. not their maximum likelihood values), we use MCMC sampling to infer *ŝ* and *ê* given the estimated rate *q_ML_* (Fig. 2B). The MCMC samples from the posterior probability

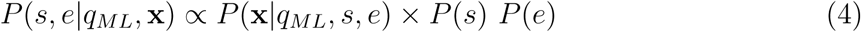

where *P* (*s*) ~ *𝒰* (*x*_1_, ∞) and *P* (*e*) ~ *𝒰* (0, *x_K_*) are uniform priors on origination and extinction times. We sample 1,000 values of *s* and *e* and use their mean as expected origination and extinction times *ŝ_q_*, and *ê_q_*. Once obtained *q̂*, *ŝ_q_*, and *ê_q_* we can calculate the likelihood of the data given the model and use it for model comparison.

Under the TPP model the expected times of origination and extinction are determined by a combination of exponential expectations with rate parameters (i.e. preservation rates) **q** = {*q*_1_*,…, q_S_*}, truncated at the boundaries of each of *S* time windows (Fig. 2C). For any given preservation rate *q*, we use numerical integration to approximate the resulting distribution and obtain expected values for the times of origination and extinction (*ŝ, ê*). We use maximum likelihood to optimize the vector of preservation rates.

The likelihood of a dataset encompassing multiple taxa, under any preservation model, is the product of the individual likelihood of each lineage (Silvestro et al., 2014b). For the purpose of model testing between HPP, NHPP and TPP models, we assume that the preservation rates are constant across lineages and therefore optimize a single parameter *q* (or vector of parameters **q** under the TPP model) to obtain the maximum likelihood of the data. We then calculate the fit of each model using the Akaike Information Criterion corrected for sample size (AICc), based on the number of analyzed lineages (Burnham and Anderson, 2002). We consider this test as a useful tool to choose between qualitatively different preservation processes (HPP, NHPP and TPP) and advise researchers to always couple the best-fitting Poisson process with the Gamma model in empirical analyses. The risk that the Gamma model represents an overparameterization of the preservation process is minimal, because the Gamma model only adds a single parameter to incorporate any potential amount of rate heterogeneity across clades (Silvestro et al., 2014b). Additionally, virtually all empirical datasets we have analyzed so far indicated very high levels of rate variation across clades (see also Results).

### AICc thresholds and testing

We used simulated data to assess the performance of our likelihood test for preservation models. We simulated 1,000 datasets of fossil occurrences under each of three models HPP, NHPP, TPP. Each simulation included 100 lineages with lifespan determined by a randomly sampled extinction rate *μ* ~ *𝒰* [0.05, 0.5], reflecting a realistic range of extinction rates (e.g. Pimiento et al., 2017). Thus, for the properties of the birth-death process (Kendall, 1948) the distribution of lifespans followed an exponential distribution with mean 1*/μ*. Fossil occurrences were then simulated based on each Poisson process with a rate *q* randomly drawn from *𝒰* [0.05, 3.5]. The rate *q* represented the mean preservation rate for each lineage in NHPP simulations (Silvestro et al., 2014b). In TPP simulations we simulated one shift in preservation rate occurring at half time between the origination time of the oldest lineage and the most recent extinction time. The preservation rate after the shift was then set to 5 *× q*.

Although singletons (i.e. lineages represented by a single fossil occurrence) can be analyzed and are usually included in PyRate analyses, they should be removed when the aim is comparing the fit of different preservation models. While singletons contribute to the correct inference of preservation rates in an analysis aimed at parameter estimation, at least one waiting time between occurrences is needed when testing among preservation models. Singletons are therefore removed automatically from the data when using the model testing function implemented in PyRate. Thus, before running the test on simulated data we removed all lineages with fewer than 2 occurrences. This procedure left, depending on the simulation settings, between 10 and 100 sampled lineages, providing a range of data sizes.

We used simulations to define the appropriate *δ*AICc thresholds necessary to confidently choose between preservation models. While the model yielding the smallest AIC score can be considered as best fitting (Burnham and Anderson, 2002), small differences in AICc values might be difficult to interpret and the threshold for significance is often obtained through simulations (e.g. Pennell et al., 2014; Dib et al., 2014). Additionally, verifying empirically the accuracy of model testing is especially important here since the optimization involves a combination of analytical expectations of origination and extinction times for HPP and numerical approximations for NHPP and TPP. Thus, we used the 3,000 simulations (for which the true generating model is known) as a training set and for each computed AICc scores under the three preservation models. Based on the resulting distributions of AICc scores, we determined the *δ*AICc thresholds yielding less than 5% errors and less than 1% errors in model selection. We then simulated an additional 300 datasets (100 for each preservation model) to verify the appropriateness of the thresholds (Fig. S1 –S3).

### Time-variable birth-death models

The temporal distribution of origination and extinction times of sampled lineages, estimated through the preservation process, is modeled to be the result of a time-continuous birth-death stochastic process, where lineages originate at a rate *λ* and go extinct at a rate *μ* (Kendall, 1948). PyRate implements several birth-death models, in which rates can change through time at discrete events or rate shifts (Silvestro et al., 2014b), following time-continuous variables (Lehtonen et al., 2017). The general likelihood of a birth-death process with time variable rates is derived from Keiding (1975):

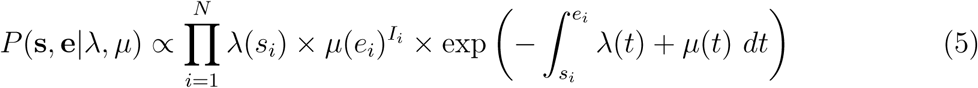

where N is the number of lineages, *λ*(*t*) is the origination rate at time *t*, *μ*(*t*) is the extinction rate at time *t* and *I_i_* is an indicator set to *I_i_* = 1 if species *i* is extinct (*e_i_ >* 0) and *I_i_* = 0 if species *i* is extant (*e_i_* = 0).

A birth-death model with rate shifts (BDS) is characterized by changes in rates of origination and extinction at shift times, while the rates are constant between shifts (Silvestro et al., 2014b). The BDS model is described by a vector of origination rates Λ = {*λ*_0_*, λ*_1_*,…, λ_J_* } delimited by times of shifts 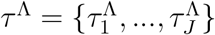 and by extinction rates *M* = {*μ*_0_*, μ*_1_*,…, μ_H_* } delimited by times of shifts 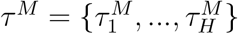, where *J* and *H* represent the number of origination and extinction rate shifts, respectively. Under this notation, origination and extinction rates are constant and equal to *λ*_0_ and *μ*_0_, respectively, when the model includes no rate shifts. The original PyRate implementation used a Bayesian algorithm, the BDMCMC (Stephens, 2000), to jointly infer the number of rate shifts (*J* and *H*), the rates between shifts (Λ*, M*) and the times of rate shift (*τ* ^Λ^*, τ ^M^*). While we showed BDMCMC to be able to correctly infer rate variation under several scenarios, it tends to be too conservative in assessing rate heterogeneity through time when the true generating process involves several rate shifts (Silvestro et al., 2014b). In the sections below we develop an alternative method to estimate birth-death models with rate shifts using the more general RJMCMC algorithm (Green, 1995), and demonstrate through simulations that it outperforms BDMCMC.

### Inferring rate variation using RJMCMC

In the RJMCMC framework the number of rate shifts is considered as an unknown variable and is estimated from the data. To this end we include two additional types of proposals: namely the *forward move* and the *backward move*, which add or remove rate shifts, respectively, thus changing the number of parameters in the birth-death model. Given that these moves are identical for both speciation and extinction rates, we use the notation Φ to denote either the speciation (Λ) or extinction (*M*) rates. We indicate the time frames identified by rate shifts with ∆ = {*δ*_0_*, δ*_1_*,…δ_K−_*_1_}. Under this notation, we set *δ_i_* = *τ_i_ − τ_i_*_+1_, where *τ* is the time of rate shift for 0 *< i ≤ K*, whereas *τ*_0_ = max(**s**) and *τ_K_*_+1_ = min(**e**) represent the maximum and minimum ages of the full birth-death process spanned by the data. A given set of time frames ∆ of length *K* is associated with a vector of rate parameters Φ = {*ϕ*_0_*, ϕ* _1_*,…, ϕ _K_*}.

The RJMCMC algorithm requires a modification in the acceptance rule of a standard MCMC in order to maintain its reversibility, while moving across models with different parameterization (Green, 1995). The general form of the acceptance probability for a *forward move* (i.e. adding a rate shift) can be written as min {1*, A*(*θ, θ′*)}, where *θ* and *θ′* are the model parameters of the current and new states, respectively and *A*(*θ, θ′*) is the product of three main terms:

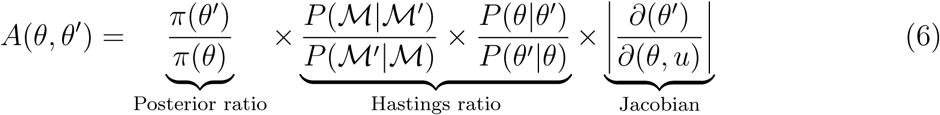

The first term is the posterior ratio, i.e. the ratio between unnormalized posterior probabilities, of the new state over the current state (where *π*(⋅) indicates the posterior as in Eq. 1). The second term, often referred to as the Hastings ratio (e.g. Heath et al., 2014), describes the ratio between the probability of going back from the new state to the current one and the probability of proposing the new state given the current one. This term includes the probability of a forward move, which generates a new model 𝓜 ′ from the current one 𝓜 by adding a rate shift and the probability of a *backward* move, which removes a rate shift. The Hastings ratio also includes the probability of proposing a new parameter state *θ′* from the current one *θ* and vice versa. Note that the new and current states will differ in the number of parameters by one additional time of rate shift and one additional rate shift. The third term is the Jacobian of the mapping function transforming the parameters of the current state into the parameters of the new state and corrects for the change in the dimensionality of the parameter space. The acceptance probability of a *backward move* (i.e. removing a rate shift) can be directly deduced from the associated *forward move*. The move from a model with parameters *θ* (with *K* rates) to a model *θ′* (with *K −* 1 rates) has the acceptance probability set to min (1*, A*(*θ′, θ*)) with

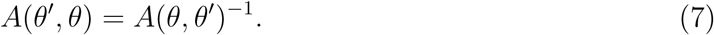

#### Probability of a reversible jump

In our implementation *forward* and *backward moves* are selected with equal probability *P* (𝓜_*K*+1_*|*𝓜_*K*_) = *P* (𝓜*_K_|*𝓜_*K*+1_) = 0.5 except for the boundary cases *K* = 1 and *K* = *K*_max_, where *K*_max_ is the maximum allowed number of rate shifts. When *K* = 1, i.e. constant rates and no rate shift, *forward moves* are proposed with probability 1, while only *backward* moves are proposed when *K* = *K*_max_. To avoid numerical issues (e.g., overflows), PyRate does not allow time windows smaller than 1 time unit (i.e. *δ >*= 1), therefore resulting in *K*_max_ = *τ_K_*_+1_ *− τ*_0_.

#### *Forward move*: adding a new rate shift

A *forward move* from model 𝓜*_K_* to 𝓜_*K*+1_ is done by splitting an existing time frame into two time frames to which new rates are assigned. We first select a time frame *δ_i_* randomly from ∆ and split it into two time frames *δ_x_, δ_y_*, by drawing a new time of rate shift *τ’* from 𝒰 (*τ_i_, τ_i_*_+1_). Since *δ_x_* + *δ_y_* = *δ_i_*, we can calculate the relative weight of the two new time frames as *w_x_* = *δ_x_/δ_i_* and *w_y_* = *δ_y_/δ_i_*. We then assign the rates *ϕ_x_* and *ϕ_y_* to the new time frames, to replace the original *ϕ_i_*. Although the new rates could be drawn from independent distributions, we choose *ϕ_x_* and *ϕ_y_* such that their weighted geometric mean equal the original rate *ϕ_i_*, which was shown to be more efficient in Poisson processes with rate shifts Green (1995). The weights are *w_x_* and *w_y_* (i.e. based on the relative size of the new time frames) and the new rates are chosen so that

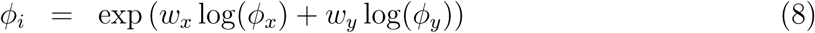

We draw a random variable *u* from a beta distribution *B*(*α, β*) that quantifies the amount of discrepancy between rates *ϕ_x_* and *ϕ_y_* by using the following equation

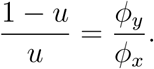

We therefore generate the new rates as:

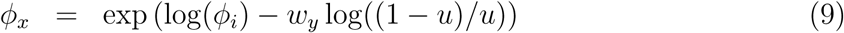

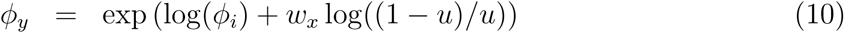

The parameters of the beta distribution are set by default to *α* = *β* = 10, yielding an expected *E*[*u*] = 0.5 with 95% of the values ranging from 0.29 to 0.71. We chose these values as they provided good convergence in our tests, although PyRate includes commands to easily tweak this and other tuning settings.

The Hastings ratio for a *forward move* 𝓜_*k*_→𝓜_*k*+1_ is computed as

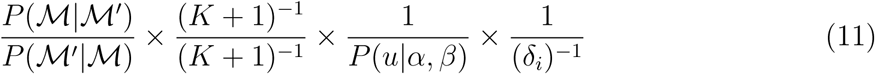

where the first ratio is based on the simple rules described above and allowing *forward* and *backward moves* with equal probabilities when 1 *< K < K*_max_. The numerator and denominator of the second ratio define the uniform probability of drawing one of the *K* rate shifts from the new model 𝓜_*K*+1_ and the uniform probability of drawing one of the *K* time frames from the current model 𝓜_*K*_, respectively (noting that a model with *K* rate shifts includes *K* + 1 time frames).The two following denominators identify the probability of drawing *u* from its distribution *β*(*α, β*) (where *P* (*u|α, β*) is based on the probability density function of a beta distribution *B*(*α, β*)) and the probability of uniformly drawing a new rate shift within time frame *δ_i_*. The Jacobian for the transformation of variables (*ϕ_i_, u*) *→* (*ϕ_x_, ϕ_y_*) (Eqn. 9) is equal to (Green, 1995):

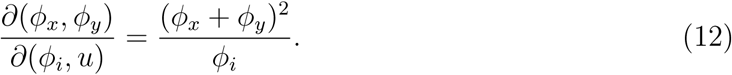

#### *Backward move*: removing an existing rate shift

A *backward move* from model 𝓜_*K*+1_ to 𝓜*_K_* is done by removing an existing rate shift and merging the two adjacent time frames and their rates. The first step is to randomly select a rate shift *j* over the *K −* 1 existing ones. The temporal placement of the rate shift is *τ_j_* and its adjacent time frames are identified as *δ_j−_*_1_ and *δ_j_*. Thus, the rates *ϕ_x_* and *ϕ_y_* are combined to obtain a new rate *ϕ_i_* based on Eq. 8.

For a *backward move* 𝓜_*K*+1_ *→* 𝓜_*K*_, the same computations are applied but the Hastings ratio and the Jacobian must be inverted as defined in Eq. (7). The value *u* must be defined using Eqs. (9) in order to compute *P* (*u|α, β*).

#### Priors on the number of shifts

Because in the RJMCMC implementation the number of origination and extinction rates (*J* and *K*, respectively) are considered as unknown variables, we assign them a prior distribution to sample them from their posterior distribution. We use a single Poisson distribution with rate parameter *r* to compute the prior probability of *J* and *K*. To reduce the subjectivity of the prior, we consider *r* itself as an unknown parameter and estimate it from the data. We assign a gamma hyper-prior, which allows us to sample *r* directly from its conjugate posterior distribution for any given *J* and *K* values:

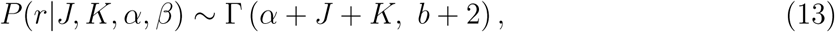

where *α* and *β* are the shape and rate parameters of the gamma hyper-prior distribution. In our simulations, we use the hyper-prior Г(*α* = 2*, β* = 1), which sets the highest prior probability to models with constant origination and extinction rates (i.e. mode = 1).

#### Marginal origination and extinction rates

To summarize the origination and extinction rates sampled by RJMCMC we marginalize them within arbitrary small (user-defined) time bins. We emphasize that this procedure does not imply that the birth-death process itself is discretized in time bins, since both the origination and extinction events are modeled within a time-continuous stochastic process. The marginal distributions of origination and extinction rates incorporate uncertainties on:

1. the true times of origination and extinction of sampled lineages, which is itself a function of the preservation process;
2. the number of rate shifts as sampled by the RJMCMC;
3. the temporal placement of the rate shifts.

We summarized the marginal rates by computing their posterior mean and 95% credible intervals (95% CI).

#### Timing of significant rate shifts

We implemented a function to assess the timing of significant rate changes based on the RJMCMC posterior samples. To this aim, we compute the frequency of sampling a rate shift (using arbitrarily small time bins) and plot them against time to assess when rate shifts are more likely to have occurred. To assess whether the frequency of a rate shift significantly exceeds the prior expectation, we run an MCMC simulation where the number and times of rate shifts are purely sampled from their respective priors, i.e. a uniform distribution on the times of shift and Poisson distributions on the number of speciation and extinction rates with a gamma prior assigned to its hyper-parameter *r* (see paragraph above).From the samples obtained from the simulation, we compute the prior probability of a rate shift at any given time, based on the user-specified size of the bins.

We then compute the posterior sampling frequencies corresponding to significant statistical support based on the standard log Bayes factors thresholds (so that 2 log *BF* = 2 and 6, for positive and strong support, respectively) (Kass and Raftery, 1995).

Given the two alternative hypotheses (presence of absence of a shift in a bin), we can define the Bayes factor as the the posterior odds divided by the prior odds (Kass and Raftery, 1995):

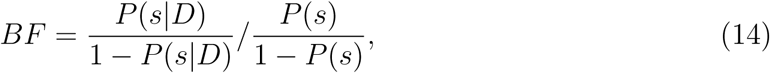

where *P* (*s|D*) is the posterior probability of a rate shift, *P* (*s*) is its prior probability. After solving the equation for the posterior term, we obtain that the posterior probability corresponding to a 2 log *BF* = *x* is

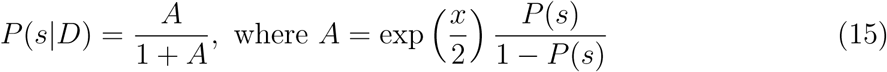

We implemented these calculations directly into a single function that generates plots of marginal origination and extinction rates through time and posterior frequencies of rate shifts through time with dashed lines indicating positive and strong statistical support based on Bayes factors (i.e. 2 log *BF* = 2 and 6, respectively; Kass and Raftery, 1995).

#### Simulations

We tested the new RJMCMC algorithm on simulated datasets and compared its performance with that of the BDMCMC algorithm previously implemented in PyRate. We simulated fossil datasets under three different birth-death scenarios:

1. Constant origination and extinction rates set to 0.15 and 0.07, respectively, with root age set to 45 Ma.
2. Time-variable birth-death model with 2 rate shifts in origination and 2 rate shifts in extinction. The time of origin was set to 35 with origination rate shifts at 20 and 10 Ma and extinction rate shifts at 15 and 10 Ma. Origination rates decrease across time windows (Λ = {0.4, 0.1, 0.01}), whereas extinction rates peaked between 15 and 10 Ma (*M* = {0.05, 0.3, 0.01}).
3. Time-variable birth-death model with 4 rate shifts in origination (at 30, 18, 15, 7 Ma) and 4 rate shifts in extinction (at 25, 22, 17, 2). Origin time was set to 45 Ma, and the rates between shifts were: Λ = {0.3, 0.07, 0.6, 0.05, 0.3} and *M* = {0.02, 0.6, 0.05, 0.2, 0.5}.

We simulated 100 datasets under each scenario assuming a homogeneous Poisson process of preservation with rate drawn from a uniform distribution *q* ~ *𝒰* [0.5, 1.5]. To avoid extremely small or large datasets, we constrained the simulations to yield between 150 and 250 lineages. We analyzed each dataset using both BDMCMC and RJMCMC, running for each algorithm 2,000,000 MCMC iterations, sampling every 1,000 iterations.

We assessed the performance of the BDMCMC and RJMCMC algorithms by quantifying their ability to infer the correct number of rate shifts and the accuracy and precision of the origination and extinction rates, marginalized within 1 Myr time bins. We computed the posterior probability of models with different numbers of rate shifts based on their sampling frequencies and compared them with the true values used to simulate the data. To quantify the accuracy of rate estimates, we used the posterior mean of the marginal rates at different times and calculated the mean absolute percentage error (MAPE), i.e. the absolute percentage error between the estimated rate (*r_est_*) and the true rate (*r_true_*), computed as (*|r_est_ − r_true_|*)*/r_true_*, averaged across rates and among simulations. We also summarized the precision of the rate estimates in terms of size of the 95% CI relative to the mean rate, again averaged across rates and among simulations.

### FastPyRateC: A new C++ library for PyRate

Because of the large number of parameters estimated in a typical PyRate analysis and due to the inherent iterative nature of MCMC algorithms, the analyses of large fossil datasets (e.g. hundreds or thousands of lineages) can be very time consuming. We therefore developed a Python module named FastPyRateC to boost the performance of the analysis. This module consists of a SWIG (http://www.swig.org/) wrapper to a fast C++ implementations of PyRate core functions such as the main likelihood functions (e.g. preservation models and most available birth-death models). This module is pre-compiled for the main operating systems (see Software availability) and can be easily compiled using a Python installation script and requires a single external dependency, the C++ boost library (http://www.boost.org/).

We assessed the improvement in performance by running analyses on three datasets of 50, 150, and 300 lineages (with 543, 1368, and 2736 fossil occurrences, respectively). We ran 100,000 RJMCMC iterations under the HPP, NHPP, and TPP models coupled with the Gamma model of rate heterogeneity among lineages. Analyses were run on a Macintosh computer with a 3.1 GHz Intel Core i7 processor. We ran with and without the FastPyRateC library to compute the speed-up achieved by the C++ library and estimate the time necessary to run the default 10M iterations, which are the default number of iterations in PyRate.

### Empirical case study

We demonstrate the new PyRate implementation by analyzing genus-level fossil occurrences of marine mammals recently compiled by Pimiento et al. (2017). The data included 535 genera, 73 of which are extant, and 4,740 occurrences spanning from the Eocene to the recent. Since the dating of most fossil occurrences is given as a temporal range, we resampled the age of each occurrence uniformly from their range and produced 10 randomized input files (as in Silvestro et al., 2014b). We then repeated all analyses on each replicate and combined the results to incorporate dating uncertainties in our estimates.

First of all, we ran the a model test to choose the most appropriate preservation model. We tested the HPP and NHPP models as well as a TPP model with rate shifts set at the boundaries between epochs in the Cenozoic. We therefore ran the subsequent analyses using the best fitting preservation model and added the Gamma option to allow for rate heterogeneity across lineages. We assumed a birth-death process with rate shifts and used the RJMCMC algorithm to determine the number and temporal placement of the shifts and the origination and extinction rates through time. After running 50 million iterations, sampling every 10,000 iterations, we combined samples of the 10 randomized datasets to infer the number of rate shifts and plot origination and extinction rates through time. The complete list of commands utilized for the empirical analyses presented here is available as Supplementary Information.

### Additional features

We incorporated several new or improved utility functions in the updated PyRate. For example, the output of RJMCMC can be processed with a single command to obtain plots of origination and extinction rates through time (posterior mean and 95% credible intervals) and estimated times of rate shift. The command also runs an MCMC simulation in the background to compute Bayes factors as described above, to determine which periods of times include a statistically significant rate shift. We also included functions to plot the number of sampled lineages through time, based on the times of origination and extinction inferred using PyRate.

Finally, we implemented a new algorithm to help researchers cleaning fossil occurrence datasets. Working with fossil occurrences often requires expert taxonomic assessment of species or genera to verify that the taxonomy is as consistent as possible within a dataset. Although such an assessment cannot be fully automatized, some data-cleaning steps can be performed in a more efficient way. One problem we have often experienced is that occurrences that are identified as belonging to one species, may be assigned slightly different Latin names (depending on the author or database). This might be due to typos or to slight variations in spelling, especially when looking at occurrences from different online databases, such as The Paleobiology Database (https://paleobiodb.org), the NOW database (http://www.helsinki.fi/science/now/), or Miomap (http://www.ucmp.berkeley.edu/miomap/). Examples of this are *Amblonyx cinerea* vs *Amblonyx cinereus* or *Felis libyca* vs *Felis lybica*. The presence of typos and spelling variation in species names can artificially inflate the number of lineages analyzed, therefore biasing the results. However, manually identifying these spelling issues can be extremely difficult and time consuming when dealing with thousands of occurrences.

We implemented, as a utility function in PyRate, a machine-learning algorithm that classifies species names (genus + species epithet) and identifies groups of names that only differ by typos or small spelling differences. We designed the algorithm specifically to deal with Latin names applying different scores to quantify differences between strings, based on common variations in Latin nomenclature (e.g. gender differences: *antiquus* vs *antiquum*). The output of this algorithm is a list of species names that are likely to represent variations of the same taxonomic entity, after which it is up to the scientist to decide if the names indeed belong to the same species and which name should be used in the final dataset. We emphasize that the algorithm does not check for synonyms (for which a look-up table would be needed), but only identifies spelling variations.

We tested this algorithm on a large fossil dataset that combined all mammalian occurrences identified to a species level retrieved from PBDB (accessed on Feb 9, 2018) and from NOW (accessed on May 9, 2017). The combined dataset included 106,937 occurrences and 19,231 unique species names.

## Results

### Testing among preservation models

The maximum likelihood test implemented to distinguish among alternative preservation processes provides a reliable tool to infer the correct model. Extensive simulations show that different *δ*AIC thresholds can be applied for different competing models. For instance if the best model (smallest AIC) is obtained for NHPP, we can reject the HPP model as a valid alternative only if *AIC_HPP_ − AIC_NHPP_ >* 3.8 (for a 5% error tolerance) or *AIC_HPP_ − AIC_NHPP_ >* 8 (for a 1% error tolerance). However, the TPP model can be confidently rejected simply based on *AIC_TPP_ − AIC_NHPP_ >* 0. The full set of thresholds derived from our simulations is given in Table 1 and incorporated in the model-test as implemented in PyRate 2.0.

**Table 1:** Thresholds for *δ*AIC estimated by simulations to test between different preservation models. Depending on the selected best model (i.e. the one with the lowest AIC score), different thresholds are applied to determine whether the model is significantly better than the alternatives (*P <* 0.05). Values in parentheses show the thresholds estimated for *P <* 0.01. Cases in which *δ*AIC values do not exceed the thresholds provided here, indicate that the evidence in the data is not sufficient to confidently choose among preservation models.

| Best model | $\delta\text{AIC}$ thresholds | | |
| --- | --- | --- | --- |
|  | HPP | NHPP | TPP |
| HPP | - | 6.4 (17.4) | 0 (0) |
| NHPP | 3.8 (8) | - | 0 (2.4) |
| TPP | 3.2 (6.8) | 10.6 (23.3) | - |

Our simulations show that the ability to statistically distinguish between preservation models (computed as *δ*AIC scores) generally increases with the size of the dataset, i.e. number of lineages and number of occurrences (Fig. SS1–SS3). Increasing preservation rates also yield stronger support for the correct model. Additionally, there is an effect of the extinction rate, whereby lower extinction rates are associated with better differentiation between preservation models. This effect is likely linked with the increased mean longevity of lineages, which therefore tend to accumulate more occurrences.

### Performance of RJMCMC compared with BDMCMC

The RJMCMC algorithm outperformed the BDMCMC alternative in most simulations (Table 2). The RJMCMC method identified the correct number of shifts in origination rates in 88% of the simulations. In comparison, the BDMCMC method identified correct model of origination in 52% of the simulations. This value is mostly driven by a consistent underestimation of rate heterogeneity in simulation scenarios 2 and 3. The RJMCMC analyses identified the correct model of extinction in 67% of the simulations. We note that the correct number of shifts in extinction rates was found in 99% of the simulations under scenarios 1 and 2, whereas under scenario 3 the algorithm consistently inferred four rates instead of five, suggesting that one of the rate shifts did not leave a significant signature on the simulated fossil data. The BDMCMC analyses correctly identified the absence of extinction rate shifts in scenario 1, but were substantially less accurate than RJMCMC analyses in finding the correct model in the case of rate heterogeneity (Table 2).

**Table 2:** Model testing using RJMCMC and the BDMCMC algorithms. The simulations (replicated 100 times) are based on different number of origination rates (*J*) and extinction rates (*K*): 1) *J* = 1*, K* = 1; 2) *J* = 3*, K* = 3; and 3) *J* = 5*, K* = 5. For each value of *J* and *K* we estimated the how frequently it was estimated as the best model by RJMCMC and BDMCMC across all replicates. Values in bold represent the frequencies at which the correct models were identified by the algorithms.

| n. shifts | Simulation 1 |  | Simulation 2 |  | Simulation 3 |  |
| --- | --- | --- | --- | --- | --- | --- |
|  | RJ | BD | RJ | BD | RJ | BD |
| $J = 1$ | **0.83** | **0.91** | 0 | 0 | 0 | 0 |
| $J = 2$ | 0.17 | 0.09 | 0.02 | 0.42 | 0.01 | 0.09 |
| $J = 3$ | 0 | 0 | **0.98** | **0.55** | 0.09 | 0.6 |
| $J = 4$ | 0 | 0 | 0 | 0.03 | 0.06 | 0.22 |
| $J = 5$ | 0 | 0 | 0 | 0 | **0.83** | **0.09** |
| $J = 6$ | 0 | 0 | 0 | 0 | 0.01 | 0 |
| $J = 7$ | 0 | 0 | 0 | 0 | 0 | 0 |
| $K = 1$ | **0.99** | **1** | 0 | 0 | 0 | 0.01 |
| $K = 2$ | 0.01 | 0 | 0 | 0.3 | 0.09 | 0.7 |
| $K = 3$ | 0 | 0 | **0.99** | **0.13** | 0.23 | 0.16 |
| $K = 4$ | 0 | 0 | 0.01 | 0.56 | 0.65 | 0.13 |
| $K = 5$ | 0 | 0 | 0 | 0 | **0.03** | **0** |
| $K = 6$ | 0 | 0 | 0 | 0 | 0 | 0 |
| $K = 7$ | 0 | 0 | 0 | 0 | 0 | 0 |

The marginal rates of origination and extinction were estimated with high accuracy by both BDMCMC and RJMCMC under scenario 1 (constant rates), with a MAPE around 0.08 to 0.15 (Table 3, Fig. SS4). In contrast, simulations based on time-variable origination and extinction rates show that RJMCMC estimates are substantially more accurate than those yielded by BDMCMC (Fig. 3; SS5). For instance for scenario 2, RJMCMC estimates marginal rates with an average MAPE of around 0.30, one order of magnitude lower than the MAPE ranging from 1.83 to 2.52 under BDMCMC. These results reflect the better ability of RJMCMC to recover the correct birth-death model, in terms of number of rate shifts (Table 2).

**Figure 3:**
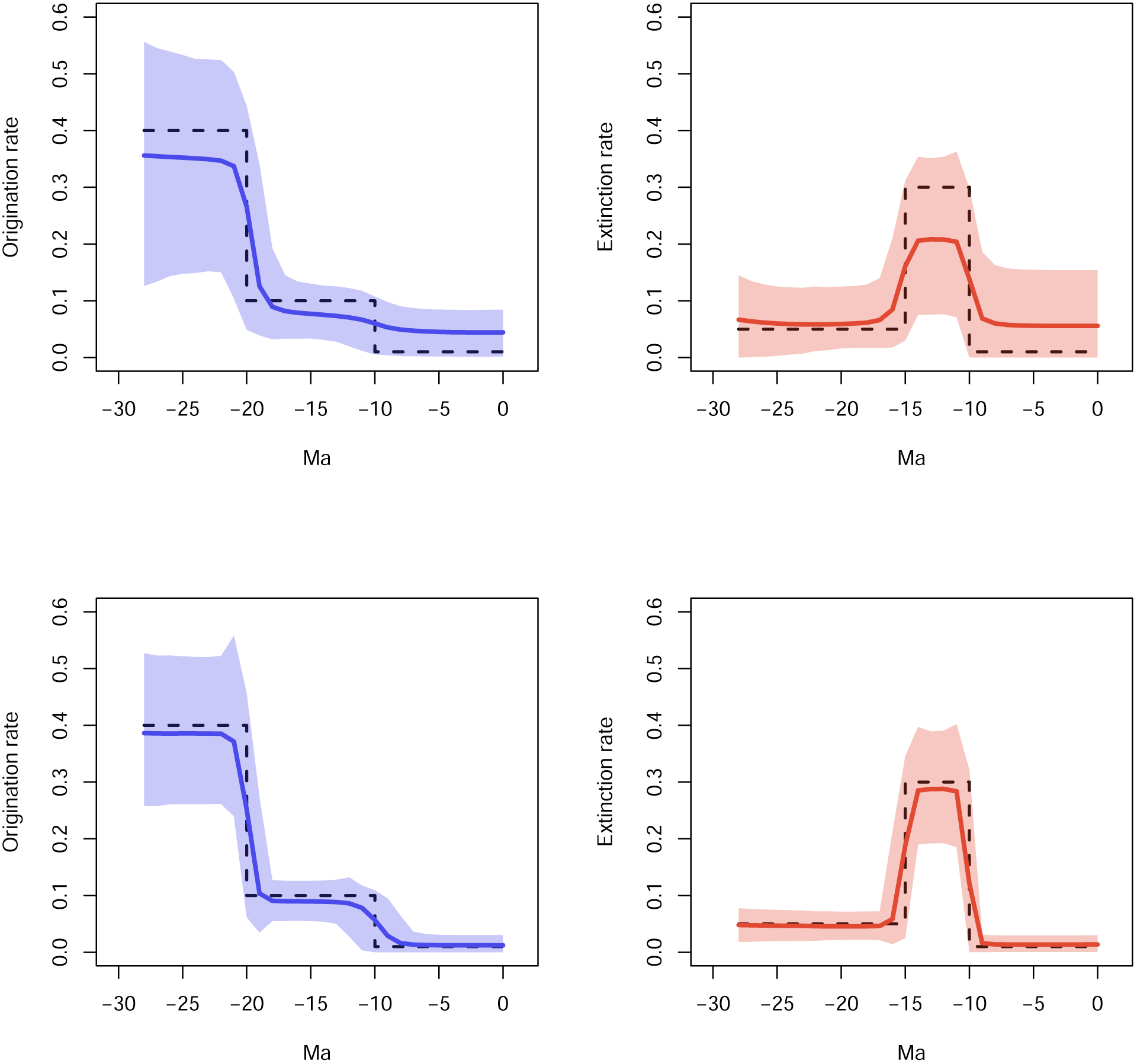
Marginal rates through time inferred for simulation scenario 2. The datasets were simulated under decreasing rates of origination (with shifts at 20 and 10 Ma) and extinction rates (with a peak at 15–10 Ma; true values are shown as dashed lines). Estimates are averaged across 100 simulations with the shaded areas showing 95% credible intervals. The top row shows the origination and extinction rates inferred using the BDMCMC algorithm, whereas the bottom row shows the results of the RJMCMC.

**Table 3:**
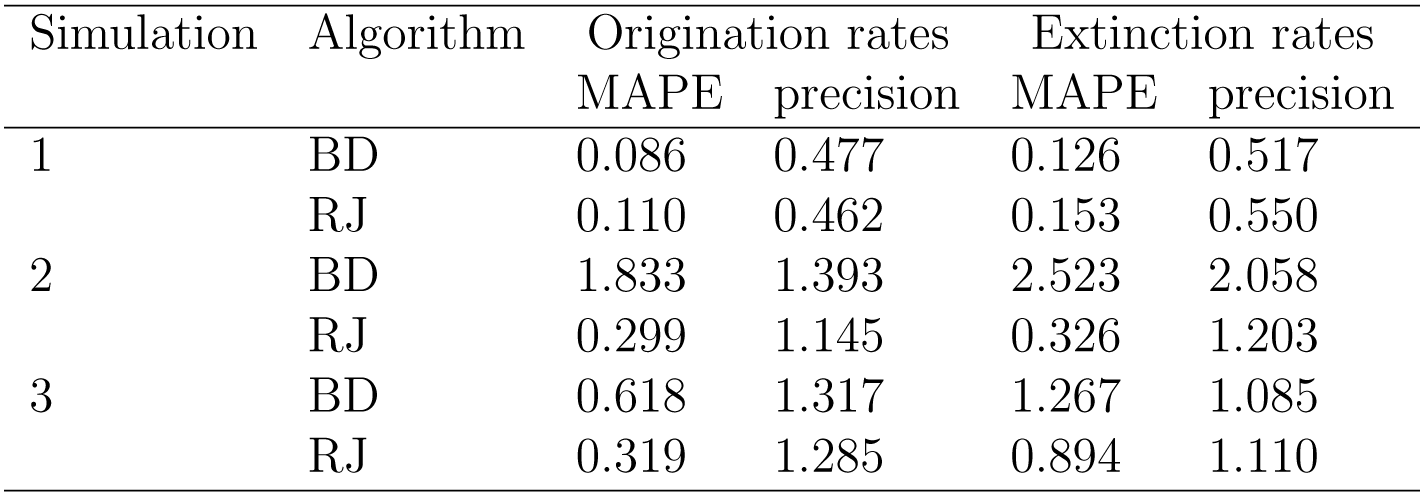
Comparison of accuracy and precision of the marginal origination and extinction rates between the new RJMCMC and the BDMCMC algorithms. Mean absolute percentage errors (MAPE) and precision are averaged across analyses of 100 simulated datasets for each simulation scenario. While the precision of rate estimates (here quantified by the relative size of the 95% credible intervals) is similar between algorithms, the RJMCMC implementation yields substantially more accurate results especially in the presence of rate heterogeneity through time.

### Performance of the FastPyRateC library

The new C++ library boosted dramatically the PyRate performance, with different levels of speed-up depending on the underlying model and the size of the dataset. In our tests the C++ version was between 5 and 8 times faster than the Python implementation when using the HPP model of preservation. Under the TPP model, the speed-up reached 26 times for a dataset of 300 taxa (Fig. 4). This performance improvement has a very significant impact on the feasibility of analyzing large dataset. For instance, an analysis of 300 taxa with TPP model, running 10 million RJMCMC iterations (default in PyRate) on a reasonably fast CPU, takes about three hours using the FastPyRateC library, whereas it takes around three days using the all-Python version. The magnitude of this performance boost becomes crucial when it comes to the analysis of large empirical datasets.The analysis of Cenozoic marine mammals presented in this study (more than 500 taxa, 50 million MCMC iterations) takes about 14 hours on a 3.1 GHz CPU, using the C++ library. In contrast, the same analysis performed using the python implementation would need more than 19 days to complete (i.e. more than 30 times longer).

**Figure 4:**
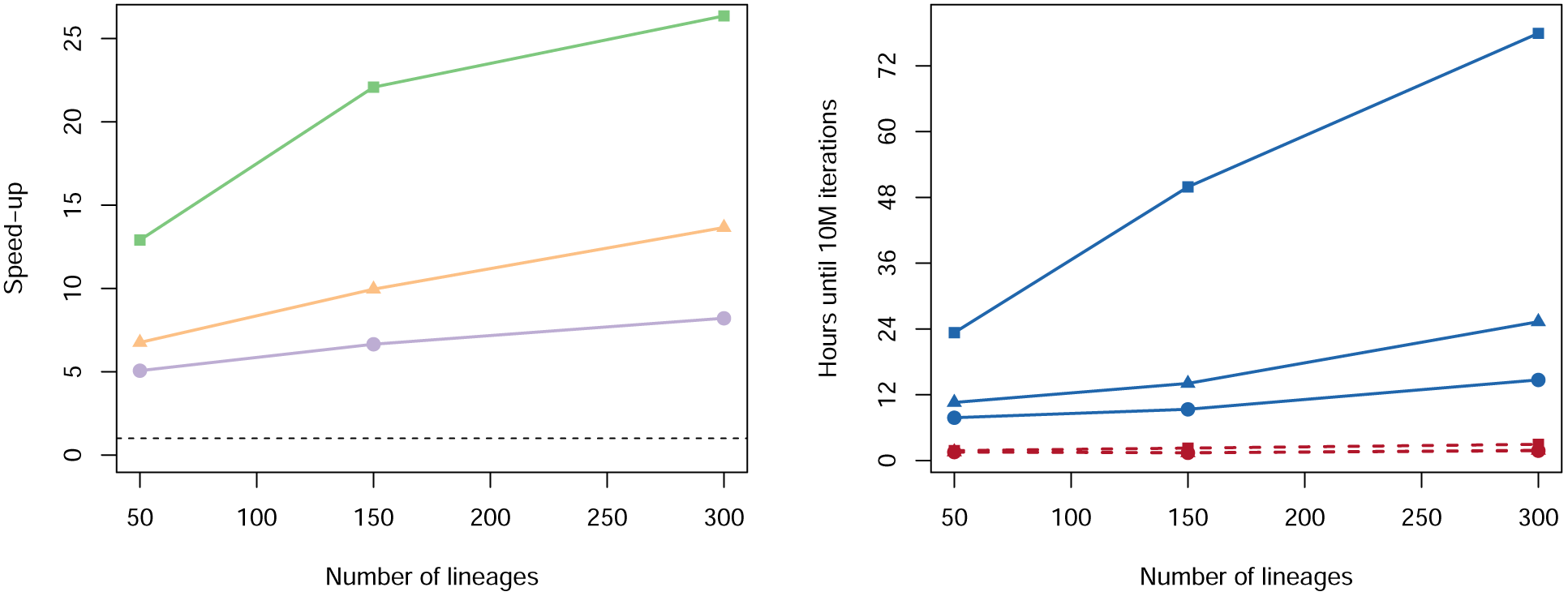
Performance comparison between the all-Python implementation of PyRate and its new version using C++ library. Comparisons are based on three datasets of 50, 150, and 300 lineages (see Methods for more details), analyzed using the RJMCMC algorithm for to infer the number and placement of rate shifts. The datasets were analyzed for 100,000 RJMCMC iterations under three preservation models: HPP (purple circles), NHPP (orange triangles), TPP (green squares).

One of the advantages of the current configuration of the FastPyRateC library (as compared to e.g. a complete re-implementation of PyRate in C++) is that the switch between Python and C++ languages happens ‘under the hood’. Thus, using or not the library does not change the way the program’s usage and PyRate automatically switches to an all-Python version if the C++ library is incompatible with the current operating system. Future program developments will be initially implemented in Python with internal functions being additionally brought to C++ to improve performance.

### Diversification dynamics of Cenozoic marine mammals

The maximum likelihood test preservation models resulted in a very strong support for the TPP model against HPP (*δ*AICc = 324.23) and against the NHPP model (*δ*AICc = 799.41). The TPP model assumed independent rates at each epoch and included 7 parameters (for Eocene, Oligocene, Miocene, Pliocene, Pleistocene, Holocene). We therefore ran the PyRate analyses using a TPP model of preservation, coupled with rate heterogeneity across lineages (Gamma model).

The estimated preservation rates showed a strong increase towards the recent. For instance, the preservation rate estimated for the Miocene was 1.15 (95% CI: 0.89–1.40), whereas in the Pliocene it was 4.06 (95% CI: 3.07–5.30), raising in the Pleistocene to 8.52 (95% CI: 6.80–10.67). Furthermore, we found evidence of strong heterogeneity of preservation across lineages, as identified by the estimated parameter *α* = 0.88 (95% CI: 0.75–1.01). This indicates that, for instance, while the average preservation rate in the Miocene was 1.15, the rate varied across lineages between 0.14 and 2.71 (median rate = 0.88).

The RJMCMC algorithm estimated a considerable amount of temporal variation in the origination and extinction rates. Constant-rate birth-death models were never sampled (i.e. null estimated posterior probability). The estimated number of rate shifts was 3 (95% CI: 2–5) for origination and 2 for extinction (95% CI: 2–5).

Origination rates (Fig. 5a) were highest in the early Eocene, indicating a rapid diversification of marine mammals, but potentially also reflecting the lack of Paleocene records in the dataset (this is also reflected in large credible intervals). After a decrease in the late Eocene, origination rates increased again during the Oligocene and early Miocene. The lowest origination rates were estimated between the late Miocene and the early Pleistocene, after which they show a mild increase. Four times of rate shift (Fig. 5b) received positive support by Bayes factors (i.e. 2log *BF >* 2) including 48–45.5, 32–29, 21–18.5, 11–15, and 1.5–1.25 Ma.

**Figure 5:**
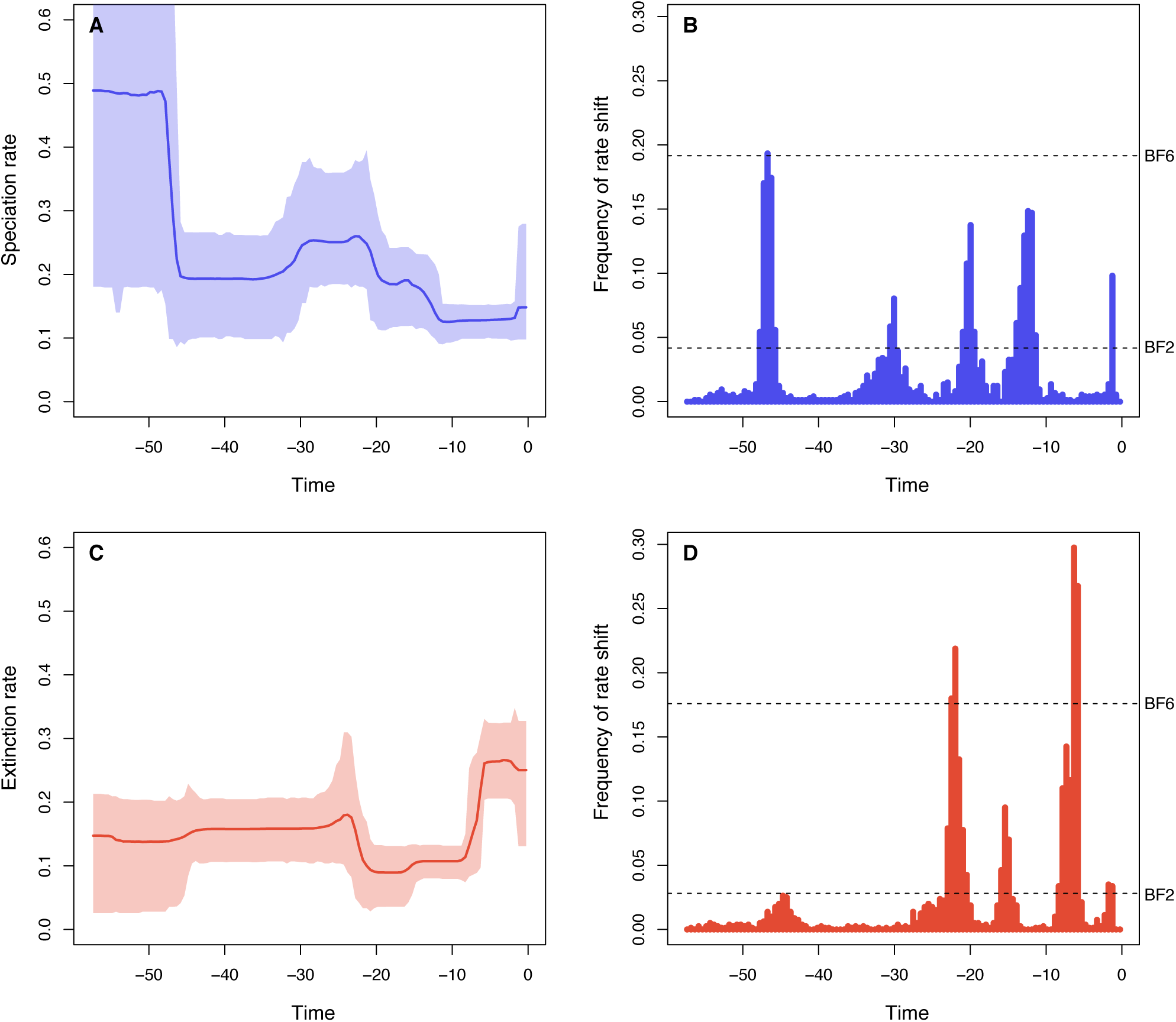
Origination and extinction rates through time in marine mammals. The dataset, obtained from Pimiento et al. (2017), comprised 535 genera and 4,740 fossil occurrences. Marginal posterior estimates of origination rates (A) and extinction rates (C) are shown together with the respective 95% credible intervals. These estimates incorporate not only parameter uncertainty, but dating uncertainties (deriving from 10 replicated analyses obtained by resampling the ages of the fossil occurrences), and uncertainties around model selection, since the RJMCMC algorithm samples the number of rate shifts from their joint posterior distribution. Plots on the right show the frequency of sampling a shift in origination (B) and extinction (D) rates within arbitrarily small time bins (here set to 0.5 Myr). Dashed lines show log Bayes factors of 2 and 6 (as inferred from MCMC simulation). Sampling frequencies exceeding these lines indicate positive and strong statistical evidence for a rate shift, respectively.

Inferred extinction rates (Fig. 5c) were stable across most of the Eocene and Oligocene and dropped in the Early Miocene. The rates increased then dramatically between the late Miocene and high levels of extinctions were inferred for the Pliocene and Pleistocene, although we estimated a mild rate decrease in the Middle Pleistocene. Bayes factors indicated strong support (i.e. 2log *BF >* 6) for rate shifts 23–21 and 6.25–5.75 Ma and positive support of shifts 16–15 and 1.25-1.75 Ma (Fig. 5d).

### Identification of spelling variations in species names

The analysis of 19,231 unique species names (global mammalian fossil occurrences from PBDB and NOW) involved the screening of 116,334,631 pairs of species names and took about 6 hours on a 3.1 GHz Intel Core i7 CPU. The function identified 174 species names as most likely (rank 0) referring to a set of 87 actual taxonomic entities. At lower similarity score (rank 1), the algorithm found 241 names which likely represent 120 actual taxonomic entities. The implemented function only flags taxa names likely representing spelling variations of the same taxonomic entity, but does not modify the original data. It is then the researcher’s task to decide which spelling is the most appropriate.

Examples of species names identified as potential variants of the same taxonomic entity (with ranks 0 or 1) included: *Deinotherium laevius* and *Deinotherium levius*, *Prosiphneus ericksoni* and *Prosiphneus eriksoni*, *Plionictis oaxacaenis* and *Plionictis oaxacaensis*, *Nannodectes gidleyi* and *Nannodectes gildeyi*. Although a detailed assessment of all these matches goes beyond the purpose of this study (but the full list of identified species names is given in Tables S1 –S4), we estimate that the fraction of false positives to be very low, with only few cases (probably fewer than 5%) identifying species names that indeed belong to different lineages, e.g. *Eomys minor Geomys minor*. The output also includes names with a lower similarity score (ranks 2–6), which almost entirely include similar names belonging to different lineages, such as *Sus arvernensis* and *Ursus arvernensis*. These results suggest that the algorithm has a very low rate of false negatives, i.e. a good power.

## Discussion

### Methodological advancements

We presented a flexible and powerful suite of quantitative methods to infer macroevolutionary processes using fossil occurrence data. These methods are part of a major update of the program PyRate and include more realistic models of preservation, new algorithms to test across models and to infer the temporal heterogeneity of origination and extinction rates.

Preservation processes are typically modeled by constant or time varying sampling probabilities (Foote, 2000; Liow and Nichols, 2010; Bapst and Hopkins, 2016), which are however constant across lineages. In PyRate, different preservation processes with constant or time-variable mean rates can be coupled with rate heterogeneity across lineages, and virtually all the empirical datasets we have analyzed so far (including the marine mammals analyzed here) support the idea that preservation varies both through time and among taxa. We demonstrated a maximum likelihood test allowing a statistical comparison among models, which facilitates an objective, data-driven, selection of the most appropriate model of fossil preservation.

We implemented a new algorithm that uses RJMCMC to estimate birth-death processes and jointly infer (in addition to the preservation parameters) the number and temporal placement of rate shifts and marginal origination and extinction rates through time. We found RJMCMC to outperform the previously implemented BDMCMC algorithm, providing more accurate rates and estimated number of shifts. The main advantages of RJMCMC are that 1) it provides marginal rates that account for uncertainties in the time and number of rate shifts, 2) it allows us to easily compute Bayes factors to assess statistically significant times of rate shift, and 3) its prior on the number of rate shifts is itself estimated from the data (unlike in BDMCMC, where it is fixed *a priori* (Silvestro et al., 2014b)), thus making the algorithm more versatile and able to adapt to different datasets.

Although the high number of parameters inferred by the PyRate model and the use of Monte Carlo sampling render the method computationally intensive, with the new C++ library we achieved a considerable speed-up (orders of magnitude). This and the ever-increasing performance of computers and clusters make PyRate a suitable method even for relatively large datasets.

### Inferring macroevolutionary rates from fossils

A large proportion of macroevolutionary research focuses on quantifying diversification process aiming to understand how biodiversity has evolved through time and space and what drives the rise and demise of clades in the tree of life (e.g. Raup and Sepkoski, 1984; Raup, 1986; Foote et al., 2007; Alroy, 2008; Quental and Marshall, 2013; Benton et al., 2014; Cantalapiedra et al., 2015; Ezard et al., 2016). The fossil record has been used to infer diversification and extinction processes for long time and arguably provides, at least for some organisms, the most informative available data for understanding macroevolutionary dynamics (Marshall, 2017).

Different approaches have been developed to this end, which typically jointly infer sampling, origination, and extinction rates (Foote, 2000; Liow and Finarelli, 2014; Alroy, 2008, 2014). PyRate is a software designed to analyze fossil data in a Bayesian framework. Its main strengths are: 1) enabling users to analyze the entire fossil occurrence record (i.e. not only first and last appearances) and all described lineages (including singletons and extant taxa) 2) incorporating parameter uncertainties using Bayesian algorithms, and 3) using explicit probabilistic model selection to infer the adequate complexity of the preservation and birth-death models based on the data. Because fossil data are often limited in size, it is essential to adequately quantify the uncertainty around each parameter estimate to avoid interpreting the results with a false sense of precision. Thus the use of a Bayesian framework is well suited for the task, providing credible intervals for each parameter rather than point estimates, and simultaneously integrating the uncertainties associated with all parameters (Gelman et al., 2013).

### Importance of model-testing in estimating origination and extinction: Comparing PyRate with other methods

Using a robust and explicit model selection framework is crucial to avoid over-parameterization and this represents one of the biggest novelties of the PyRate method, compared with other approaches. Indeed, treating origination, extinction and preservation rates in predefined time bins as independent parameters (i.e. without explicitly model-testing) is common practice in paleobiological studies of macroevolution (Foote, 2003; Liow and Finarelli, 2014; Alroy, 2015), and analogous models are available in PyRate as well (Silvestro et al., 2015b). However, this practice may generate spurious results if the amount of data is insufficient to confidently estimate all the parameters (Smiley, 2018), which is a general problem with overparameterization (Burnham and Anderson, 2002). The RJMCMC algorithm presented here and the other algorithms implemented in PyRate infer the amount of rate variation directly from the data.

Although we focused here on algorithms that simultaneously optimize the parameters and the model (RJMCMC and BDMCMC), other methods to avoid overparameterization are available in PyRate, based on the estimation of model marginal likelihoods (Silvestro et al., 2014b), Bayesian variable selection (Silvestro et al., 2015a), and Bayesian shrinkage (Silvestro et al., 2015b, 2017). Using these methods, the complexity of the model adapts to the signal provided by the data and their statistical power, so that only statistically significant rate changes are identified. This procedure also provides a formal approach to assess whether apparent rate variations are not just the result of the stochastic nature of a constant rate birth-death process.

In order to demonstrate the general importance of explicit model testing in the estimation of origination and extinction rates, we replicated some of the analyses recently presented by Smiley (2018). Smiley (2018) tested the performance of three methods, namely per capita rate method (Foote, 2000), the three-timer method (Alroy, 2008) and the capture-mark-recapture (CMR) method (Liow and Finarelli, 2014) under several preservation and diversification scenarios.

Here, we analyzed datasets simulated under constant speciation and extinction rates (set to *λ* = 0.2 and *μ* = 0.1) with low preservation rate (so that the sampling probability per lineage per Myr equals 0.3), i.e. following step-by-step the simulation settings of Smiley’s scenario “R30%”. We then generated and analyzed additional datasets following Smiley’s scenario “IncR” (where the sampling probabilities increased linearly through time from an initial 0.10 to 0.50), and scenarios “StratR” and “FreqR”, where preservation rates change over times as predicted by empirical data (based on the rock record and on North American fossil record, respectively) (Smiley, 2018). We simulated 100 datasets under each preservation scenario and analyzed them in PyRate, using the RJMCMC algorithm to infer origination and extinction rates and any evidence of rate variation and summarized the results across simulations.

PyRate correctly inferred that origination and extinction rates were constant through time under all preservation scenarios and the estimates are substantially more robust and less volatile than those from other methods which do not explicitly optimize the number of parameters in the model based on the available data (Fig. 6). The credible intervals inferred by PyRate also show that decreasing preservation rates reduce the level of confidence in origination and extinction rate estimates (Fig. 6B–D), as expected (Smiley, 2018). Although a formal comparison between the performance of PyRate and other methods is beyond the scope of this study, these results indicate that optimizing the complexity of the model based on the data is crucial to obtaining realistic estimates of diversification processes from incomplete fossil data. Based on these results, we recommend to always verify the statistical support for the number of model parameters, when inferring diversification dynamics from fossil data.

**Figure 6:**
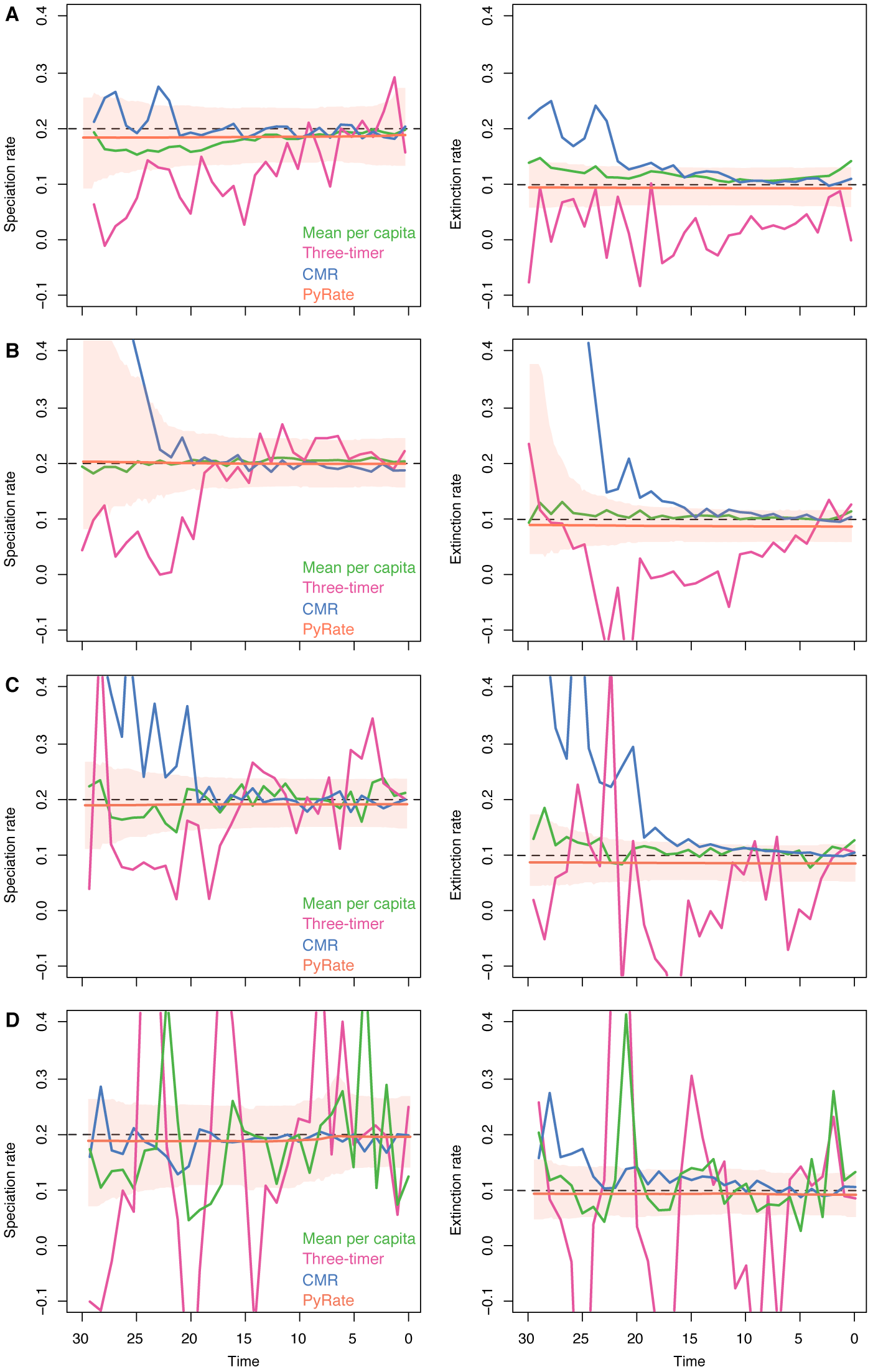
Origination and extinction rates estimated using different methods. The dashed lines indicate the true origination and extinction rates used to simulate the data. Preservation rates were constant in panel A (“R30%”), increasing through time in B (“IncR”), and varying according to empirical estimates in C and D (“stratR” and “FreqR”, respectively). See main text and Smiley (2018) for more details. Green lines show the mean per capita rates based on Foote (2000); purple lines show rates inferred using the three-timer method by Alroy (2008); blue lines indicate rates inferred using the CMR method by (Liow and Finarelli, 2014). These plots are modified from Smiley (2018). The orange lines show the posterior rate estimates inferred by PyRate using RJMCMC (summarizing results from 100 simulated datasets), with shaded areas indicating the 95% credible intervals.

## Conclusions

PyRate is an open-source project in which researchers are welcome to contribute code, ideas, and feedback through it’s Github repository. It includes numerous birth-death models for taxonomic diversification as well as several preservation models in which rates can vary through time and across lineages.The hierarchical Bayesian methods implemented in PyRate allow users to assess the statistical support of different models and to jointly infer all the parameters. Credible intervals are inferred for all model parameters (e.g. preservation, origination, and extinction rates) and can be used to quantify the level of uncertainties surrounding the estimates.

Importantly, PyRate requires a minimum number of *a priori* decisions from the user and, while each setting can be accessed through specific commands, default values and settings are set to adapt to most datasets. PyRate runs as a stand-alone command-line program and running the software does not require any knowledge of Python from the user. The program’s package also includes many utility functions that can be used to plot and summarize the results, process multiple output files, and parse large datasets to identify potential spelling variation in taxon names using a built-in machine learning classifier.

Although we focused here on diversification processes in which origination and extinction rates change through time, several other models have been implemented in PyRate enabling users to test specific hypotheses, e.g. about diversity dependent diversification with competition within and among clades (Pires et al., 2017), correlations to biotic and abiotic factors (Lehtonen et al., 2017), age-dependent and trait-dependent extinction rates (Hagen et al., 2017; Piras et al., 2018). The versatility of PyRate’s Bayesian hierarchical models enables researchers to analyze the growing amount of available fossil occurrence data and assess alternative hypotheses in a statistically robust framework.

## Software availability

All the models described in this study are implemented within the open-source package PyRate and available at: https://github.com/dsilvestro/PyRate. The program is written in Python 2.7 and R and has been tested under the major operating systems (MacOS, Windows, and several Linux distributions). A detailed command list and tutorials are available in the GitHub repository. In order to provide an easy access to the augmented performance of the FastPyRateC library, we pre-compiled modules for 64 bits versions of Windows, MacOS, and Linux and are available on the PyRate Github repository, in addition to the source code.

## Acknowledgments

All analyses were run at the High-performance Computing Center (Vital-IT) from the Swiss Institute of Bioinformatics. D.S. received funding from the Swedish Research Council (2015-04748). A.A. received funding from the Swedish Foundation for Strategic Research, the Swedish Research Council (B0569601), the Faculty of Sciences at the University of Gothenburg, the David Rockefeller Center for Latin American Studies at Harvard University, and a Wallenberg Academy Fellowship. N.S. received funding from the University of Lausanne (Switzerland) and the Swiss National Science Foundation (CR32I3-143768). X.M. received funding from the Swiss National Science Foundation (P2GEP2 178032). We thank Tara Smiley, Brianna McHorse, Marco Crotti, Torsten Hauffe, Juan L. Cantalapiedra, Oscar Inostroza, Tiago B. Quental, Mathias M. Pires, Erik Gjesfjeld for discussion and feedback on the software. We thank Etienne Orliac of the Center for Advanced Modeling Science (Switzerland) for support on the Windows version of FastPyRateC.

## Supplementary materials

### Analysis protocol for marine mammals

We list below the complete list of commands we used in the empirical analysis presented in this study. Note that all commands should be provided as a single line in a terminal (or command prompt), i.e. line breaks used below for graphical reasons should be ignored when reproducing the analyses. All datasets and input data listed below are available at https://github.com/dsilvestro/PyRate in the <u>dataPimientoEtAl2017NEE</u> directory.

#### Generate input data (in R)

Load the `pyrate utilities` script in R (the script is available in the GitHub repository) and use it to convert the tab-separated table of fossil occurrences, named”fossil occs.txt”, (from Pimiento et al., 2017) into a PyRate-formatted input file:

~~~
source(pyrate_utilities.r)
extract.ages(‘fossil_occs.txt’, replicates = 10)
~~~

This command produces a file named “`fossil occs PyRate.py`”, which can be used for analysis in Pyrate. We renamed the file to “`occs.py`” to shorten the commands below.

#### Test among preservation models (in a command-line console)

We first test between three preservation models (HPP, NHPP, TPP), where the TPP model was set to assume independent preservation rates within each geological epoch. The boundaries of the epochs are based on http://www.stratigraphy.org and given in a text file named “`epochs q.txt`”:

~~~
python PyRate.py occs.py -qShift epochs_q.txt -PPmodeltest -filter_taxa mammals.txt
~~~

This command launches the maximum likelihood algorithm and the results are printed on screen, providing the maximum likelihood values under each model, and the AICc scores that can be used for model testing (see main text). The screen output also shows which model is preferred and its level of significance compared with other models, based on the AICc thresholds derived from simulations (see main text). Note that, since the original dataset contained other marine megafauna organisms whereas here we decided to focus on mammals only, we used the command `-filter taxa mammals.txt` to provide a list of mammalian taxa that we want to include in the analysis (whereas all other lineages are dropped).

#### Run main analysis (in a command-line console)

~~~
python PyRate.py occs.py -j <rep_n> -A 4 -n 50000000 -s 10000
               -filter_taxa mammals.txt
               -qShift epochs_q.txt -mG -pP 1.5 0
~~~

where: `rep_n` is the replicate number (here ranging from 1 to 10 in ten replicated analyses), `-A 4` specifies that the RJMCMC algorithm should be used, `-n` specifies the number of iterations, `-s` specifies the sampling frequency, `-qShift` specifies that preservation is modeled by a TPP process with independent rates for each epoch, `-mG` specifies that the TPP model should be coupled by a Gamma model of rate heterogeneity across lineages, and `-pP` 1.5 0 specifies the shape and rate parameters of the gamma prior on the preservation rates. By setting the rate parameter to 0 we define the parameter as unknown, meaning that PyRate will estimate it after assigning it a hyper-prior (see main text).

This analysis produces four output files for each replicate: a summary text file with all the settings used in the analysis and three log files containing the posterior parameter values sampled by the RJMCMC. More details are provided in the online tutorial

#### Combine mcmc log files into one (excluding burnin)

PyRate includes a utility function to combine output files from different runs into one file. Assuming that all output files form the previous analyses are in the same `pyrate mcmc logs` directory, the log files are combined using:

~~~
python PyRate.py -combLog /pyrate_mcmc_logs -b 1000 -tag mcmc -resample 100
python PyRate.py -combLog /pyrate_mcmc_logs -b 1000 -tag sp_rates -resample 100
python PyRate.py -combLog /pyrate_mcmc_logs -b 1000 -tag ex_rates -resample 100
~~~

where: `−combLog /pyrate mcmc logs` provides the full path to the log files, `−b` 1000 specifies that the first 1,000 samples should be removed as burn-in, `-tag x` specifies that all files containing x in the file name should be combined, and `-resample 100` specifies that 100 random samples should be taken from each replicate and saved into the combined log files. These commands generate output files named `“combined 10mcmc.log”`, `“combined 10sp rates.log”`, and `“combined 10ex rates.log”`.

### Summarize and plot the results

The `“sp rates.log”` and `“ex rates.log”` files can be used to generate rates-through-time plots using the function:

~~~
python PyRate.py -plotRJ /pyrate_mcmc_logs -tag combined -grid_plot 0.5
~~~

where `-plotRJ /pyrate mcmc logs` specifies the full path to the log files, `-tag combined` specifies that only files containing “combined” in the file name should be plotted (by default all log files are plotted individually in a single PDF file), and `-grid plot 0.5` defines an arbitrarily small bin size used for plots and to compute Bayes factors.

This will generate an R script and a PDF file with the RTT plots showing speciation and extinction rates through time. It will also show histograms with the inferred times of rate shifts and calculate Bayes factors to help determining the time when a rate shift is supported by significant posterior probability. The histograms include two horizontal dashed lines showing the thresholds for positive evidence of a rate shift (bottom line: logBF = 2) and for strong evidence of a rate shift (top line: logBF = 6). Thus, any point in the histogram showing sampling frequencies for a rate shift exceeding the thresholds indicate a time of significant rate change.

To quantify the estimated the number of shifts we use:

~~~
python PyRate.py -mProb pyrate_mcmc_logs/combined_10mcmc_files.log
~~~

with the results (printed on screen) providing a summary of the most likely numbers of shifts in origination and extinction rates, as inferred by RJMCMC.

---

**Figure S1:**
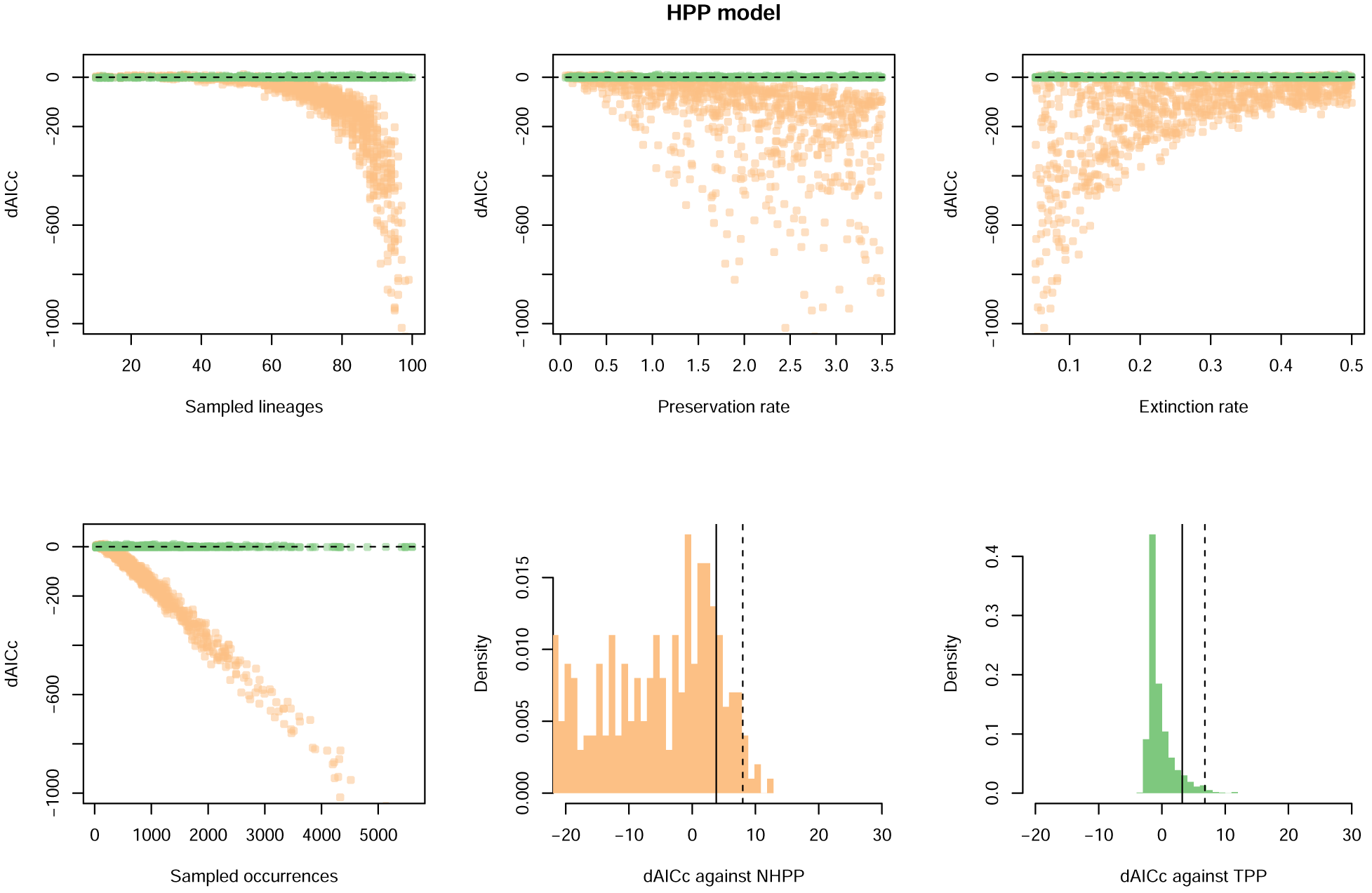
Results of model testing when the true model is HPP. Differences in AICc scores are calculated against alternative models NHPP (in orange) and TPP (in green) and plotted against several parameters used in the simulations. Scatter plots show that the ability to statistically distinguish HPP from NHPP increases with the size of the dataset, with increasing preservation rates, and with decreasing extinction rates. The two histograms (arbitrarily truncated at dAICc = −20) show the difference in AICc between HPP and the alternative models. Solid lines indicate the estimated thresholds that yield less than 5% error rate, dashed lines indicate the 1% thresholds (see main text).

**Figure S2:**
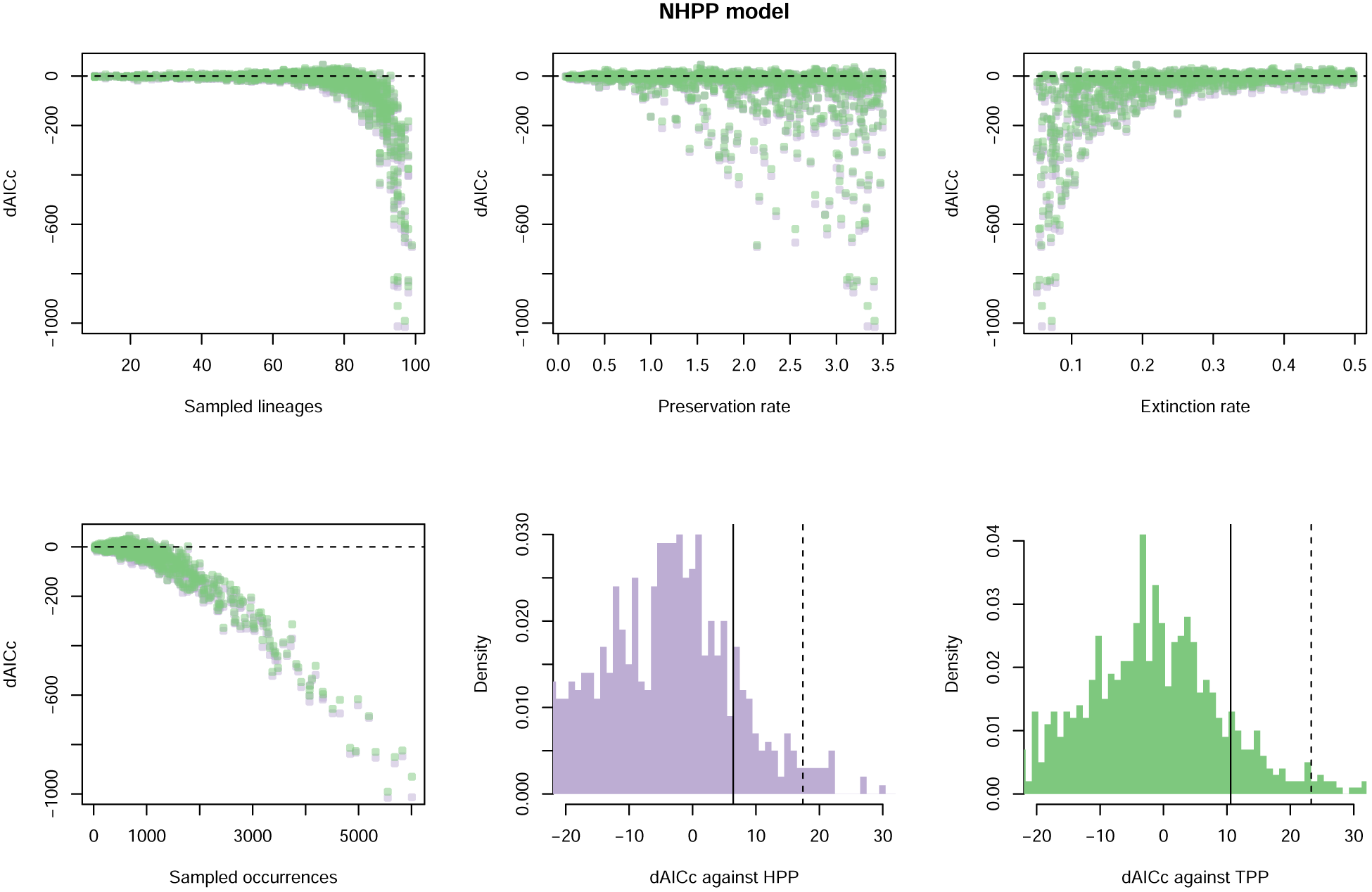
Results of model testing when the true model is NHPP. Differences in AICc scores are calculated against alternative models HPP (in purple) and TPP (in green) and plotted against several parameters used in the simulations. The two histograms (arbitrarily truncated at dAICc = −20) show the difference in AICc between NHPP and the alternative models. Solid lines indicate the estimated thresholds that yield less than 5% error rate, dashed lines indicate the 1% thresholds (see main text).

**Figure S3:**
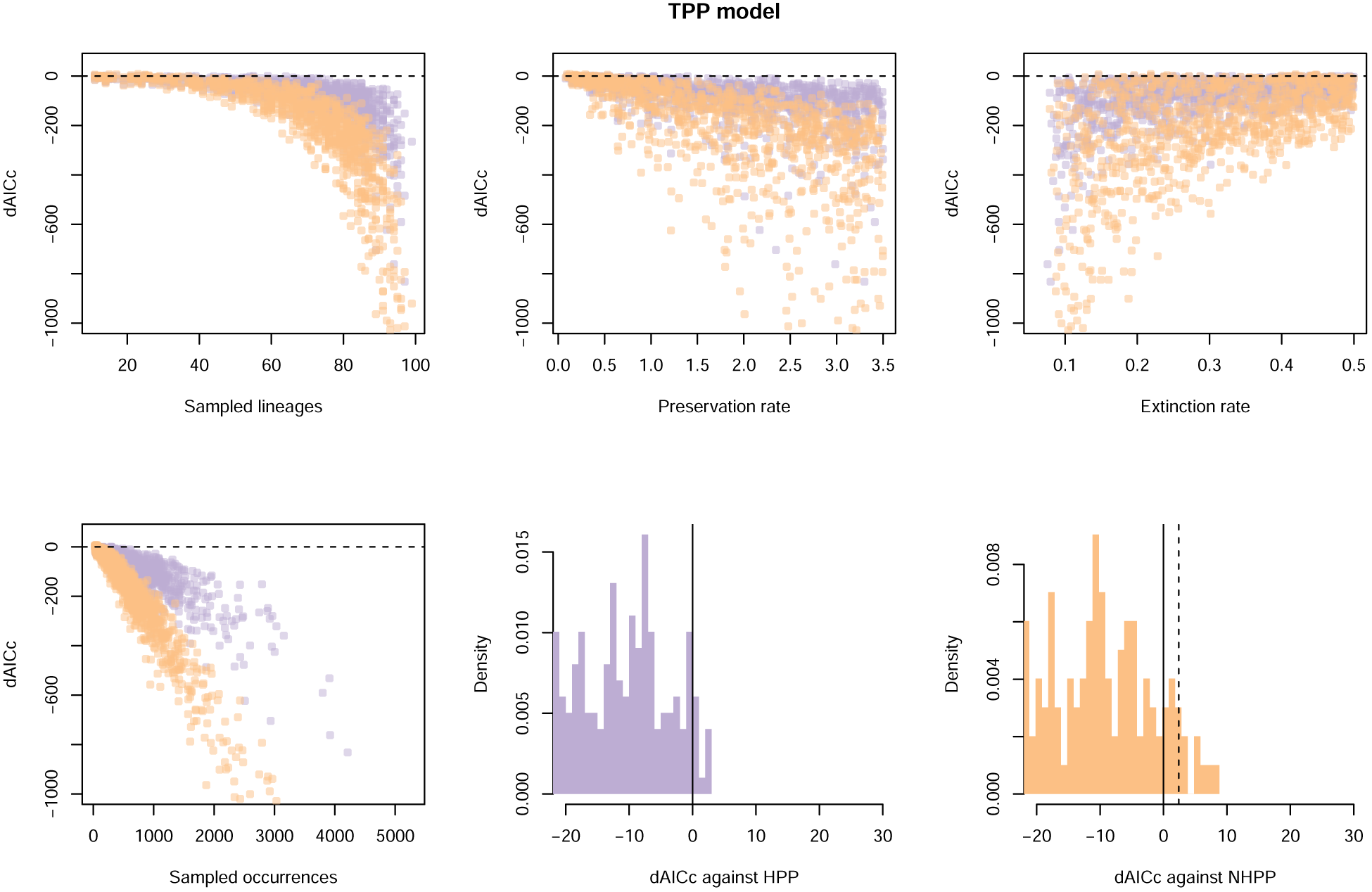
Results of model testing when the true model is TPP. Differences in AICc scores are calculated against alternative models HPP (in purple) and NHPP (in orange) and plotted against several parameters used in the simulations. The two histograms (arbitrarily truncated at dAICc = −20) show the difference in AICc between TPP and the alternative models. Solid lines indicate the estimated thresholds that yield less than 5% error rate, dashed lines indicate the 1% thresholds (see main text).

**Figure S4:**
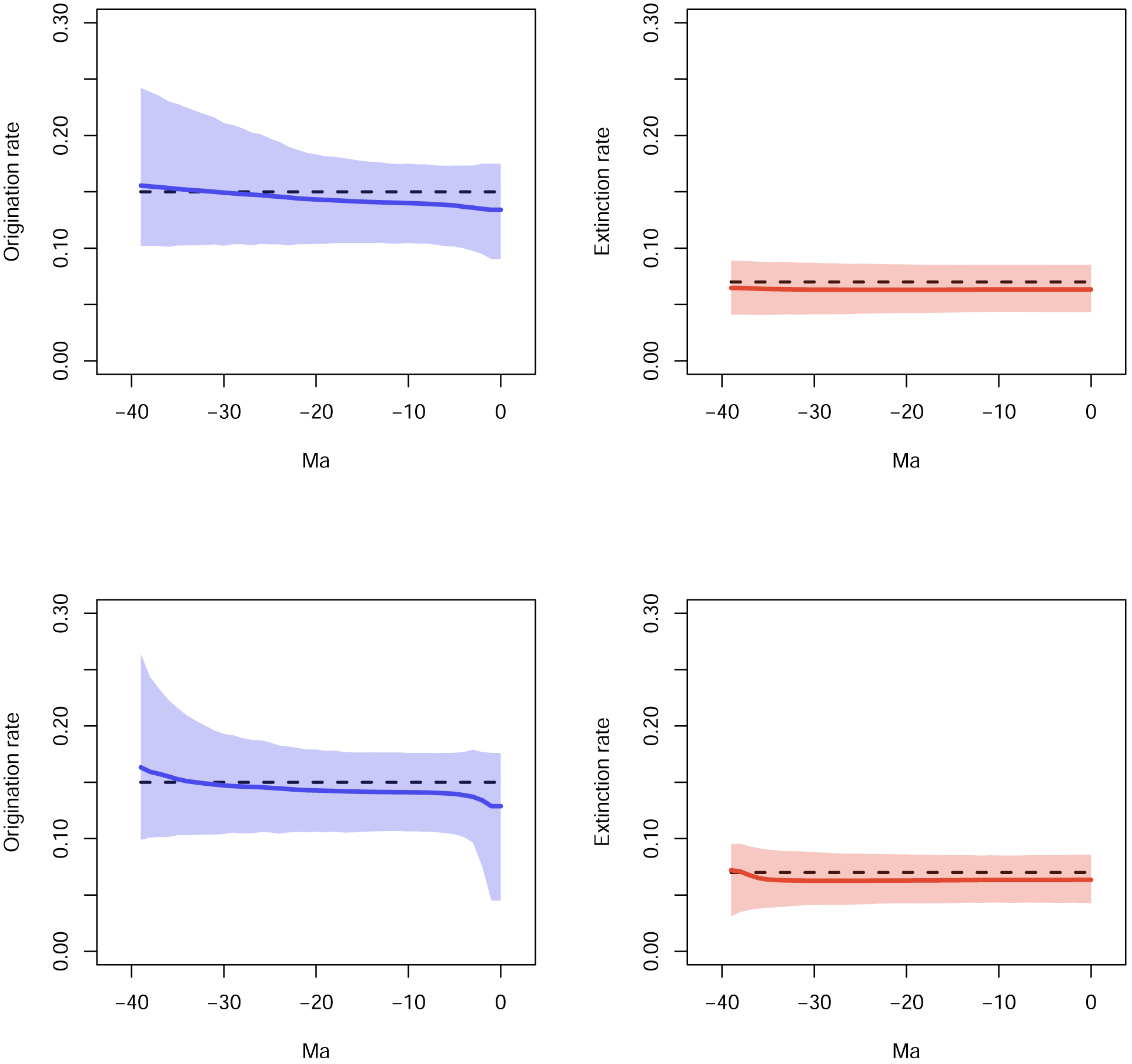
Marginal rates through time inferred for scenario 1. The dataset were simulated under constant rates origination and extinction rates (true values shown as dashed lines). Estimates are averaged across 100 simulations with the shaded areas showing 95% credible intervals. The top row shows origination and extinction rates inferred using the BDMCMC algorithm, whereas the bottom row shows the results of the RJMCMC.

**Figure S5:**
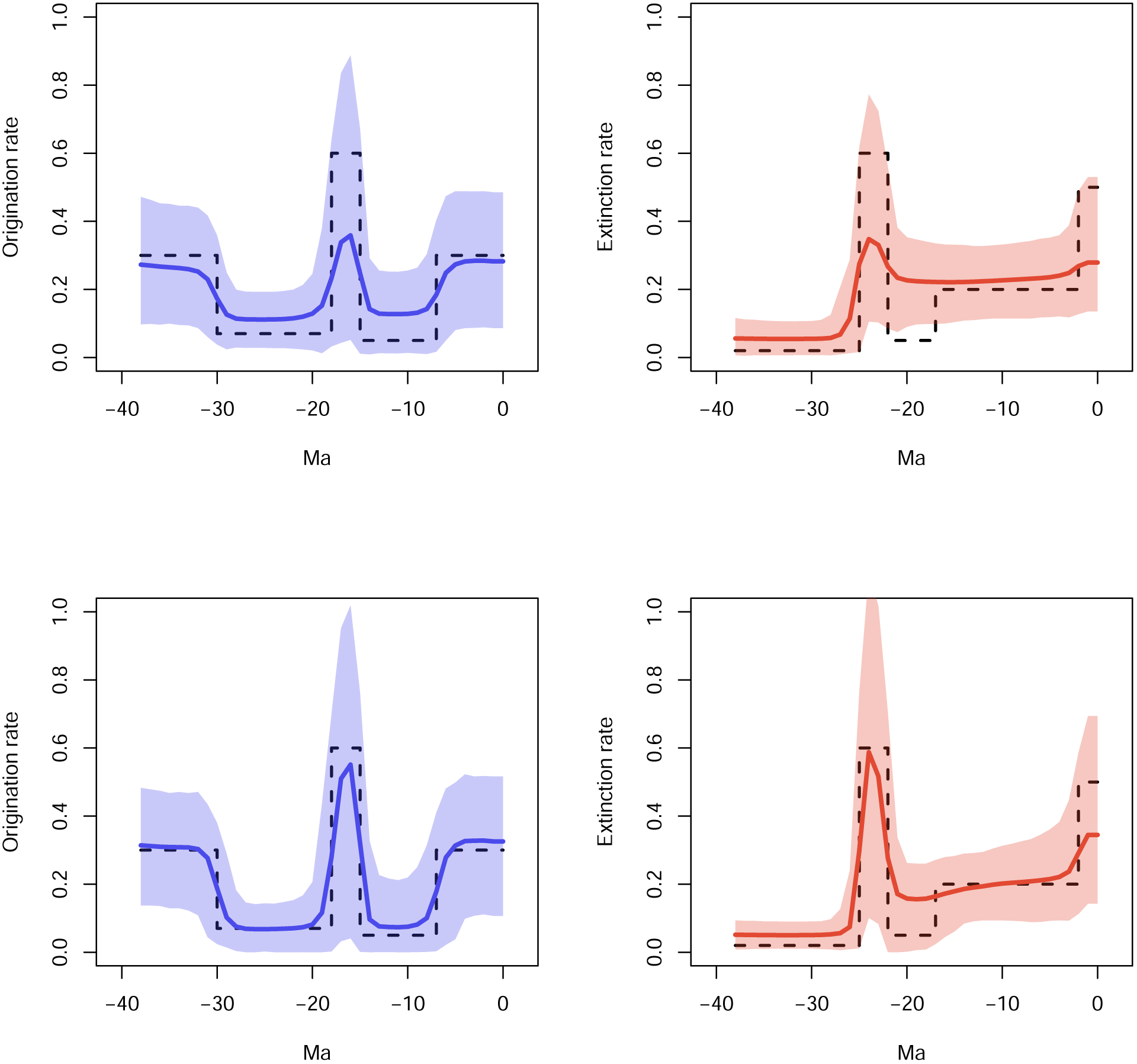
Marginal rates through time inferred for scenario 3. The dataset were simulated under variable rates origination and extinction rates (true values shown as dashed lines). Estimates are averaged across 100 simulations with the shaded areas showing 95% credible intervals. The top row shows origination and extinction rates inferred using the BDMCMC algorithm, whereas the bottom row shows the results of the RJMCMC.

**Table S1:**
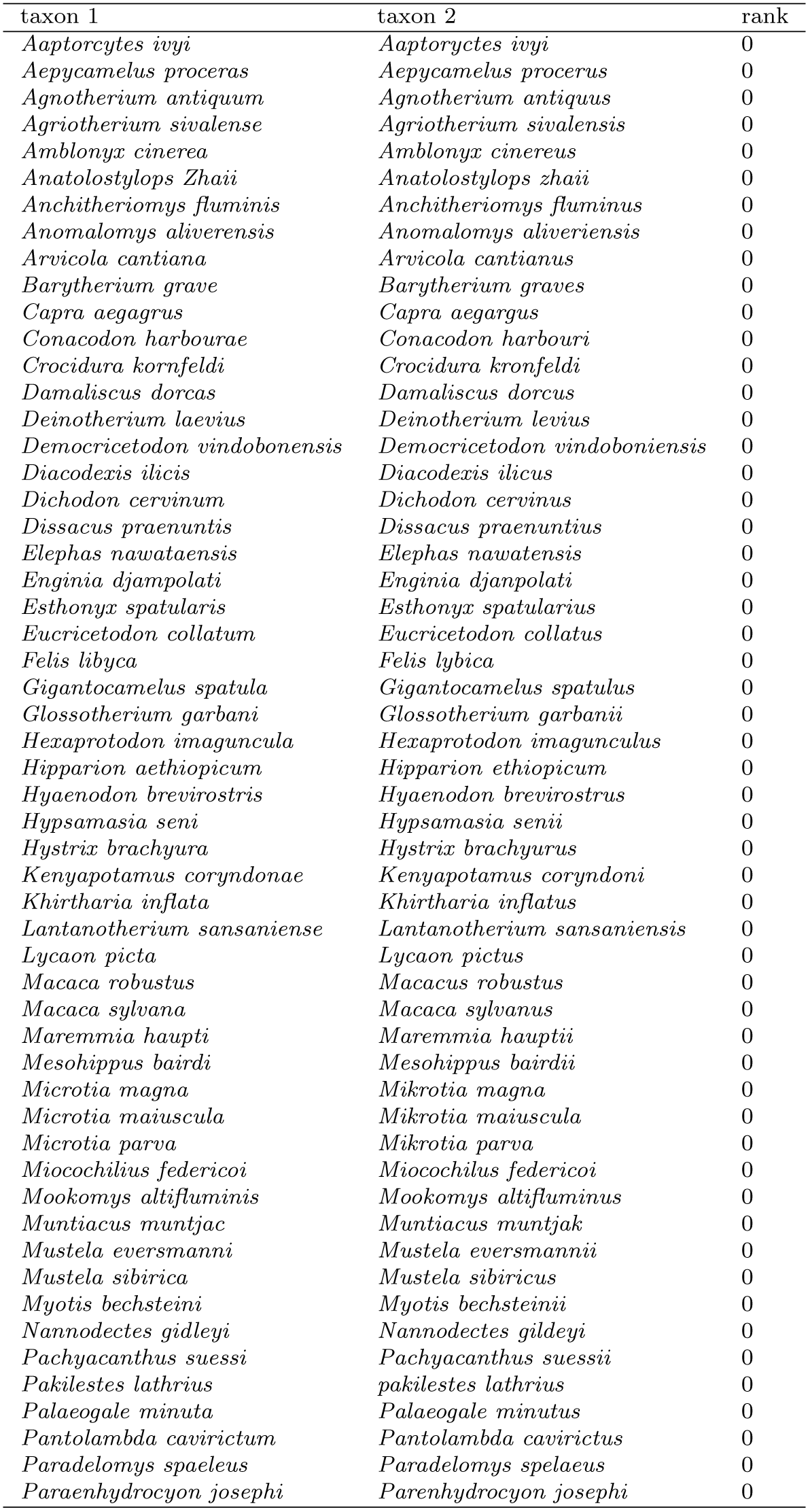
Identified variation in species name spelling. Lower rank indicate higher confidence that a pair of species names in fact refer to a single taxonomic entity. Although we report here only pairs of names ranking 0 and 1, our algorithm returns results at higher ranks as well, which are however more likely to group names with some degree of similarity, but referring to different taxa.

**Table S2:**
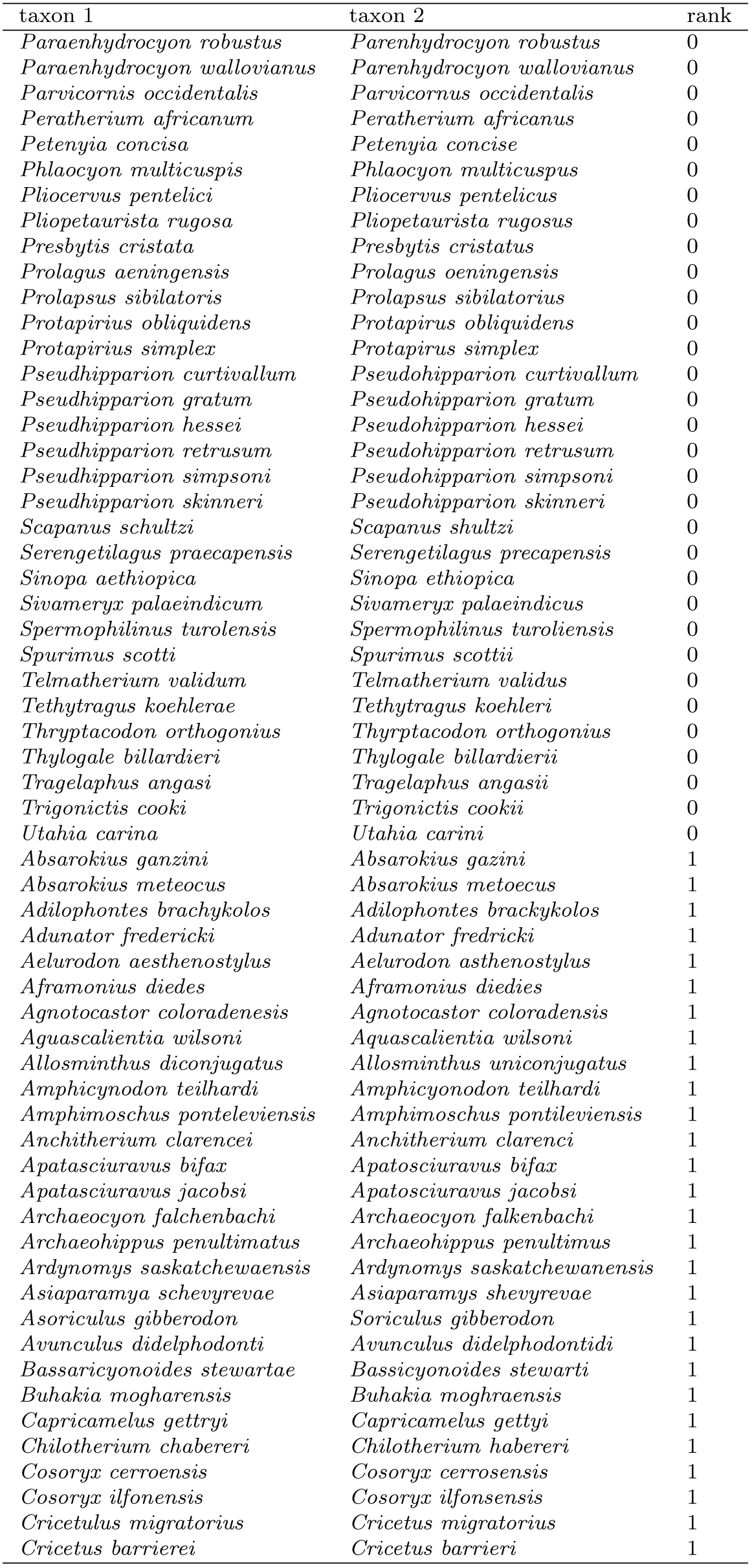
Identified variation in species name spelling - continued

**Table S3:**
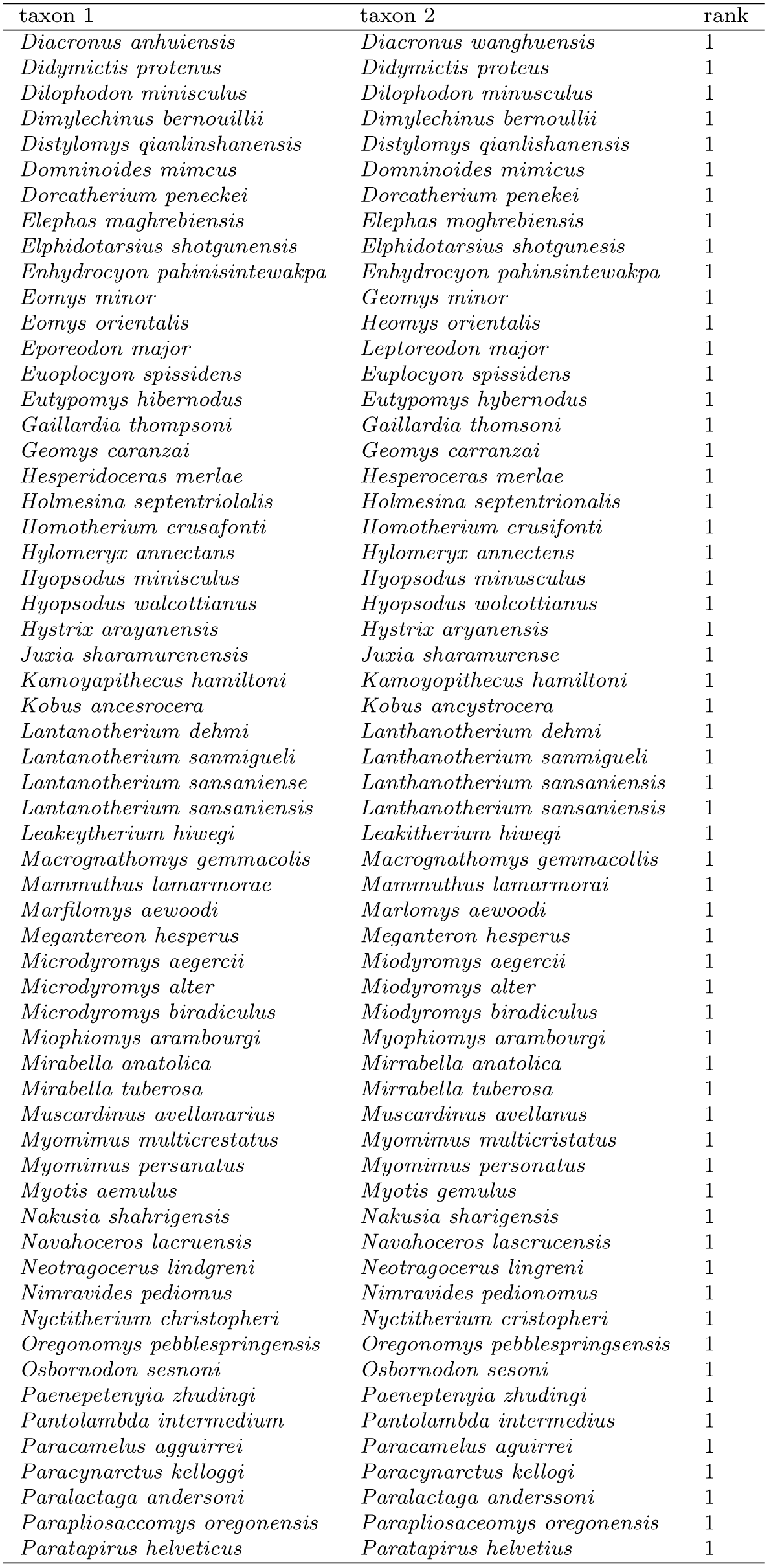
Identified variation in species name spelling - continued

**Table S4:**
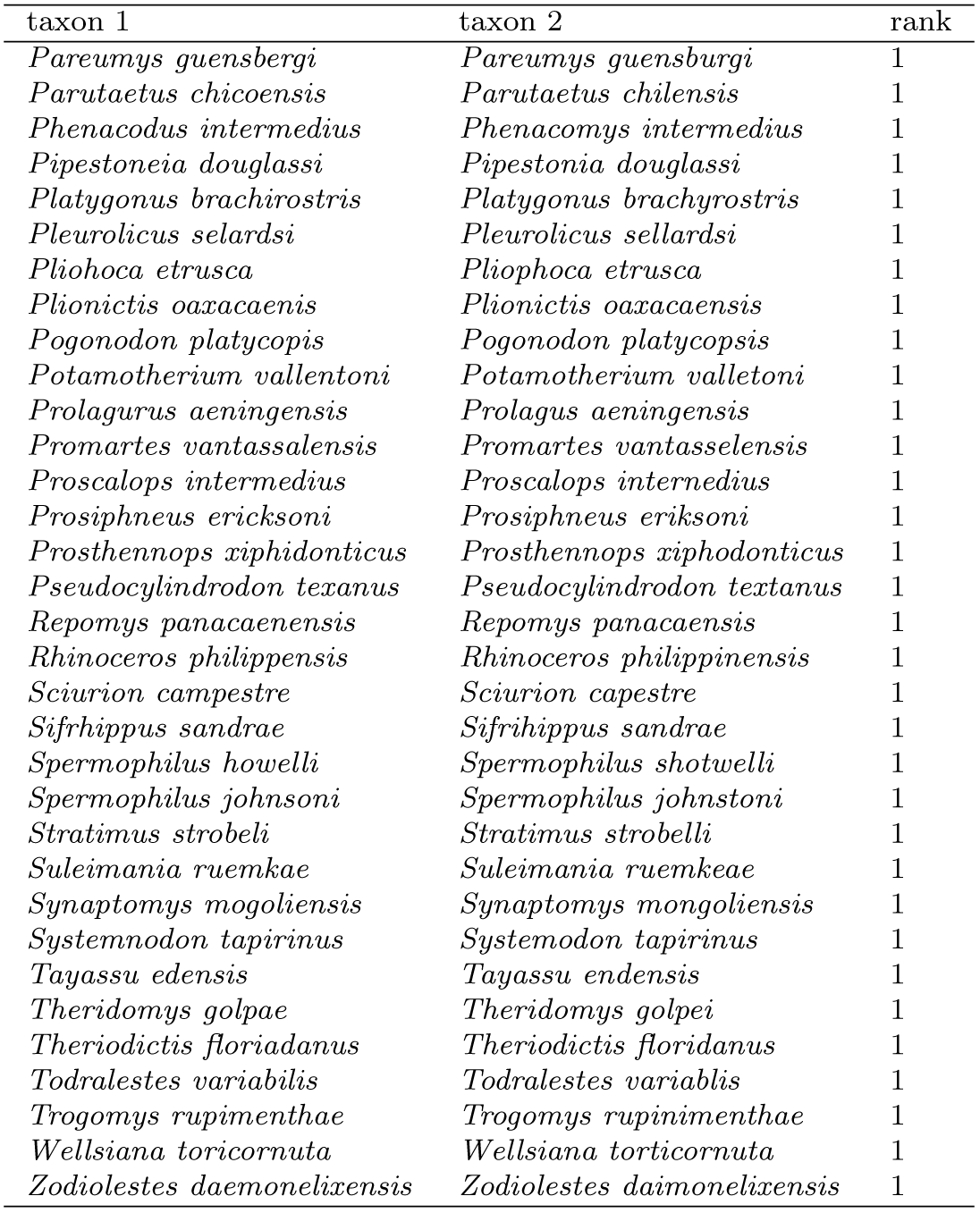
Identified variation in species name spelling - continued

